# Nuclear Irregularity as a Universal Diagnostic Tool in Solid Tumors

**DOI:** 10.1101/2025.08.12.669986

**Authors:** Francesco Hamilton, Kane Foster, Irene Ghobrial

## Abstract

As tumors develop, cancer cells accumulate diverse genomic and phenotypic alterations to meet heightened demands for energy production and biosynthesis. Loss of lamina function and perturbations in energy production are associated with pronounced aberrations in cellular morphology, particularly within nuclear architecture and the plasma membrane. Systematic analysis of nuclear morphology can reveal conserved structures across diverse cancer types, enabling disease state stratification, biomarker discovery, and potential avenues for personalizing therapy to minimize recurrence risk. To this end, this study analyzes an imaging mass cytometry (IMC) breast cancer dataset, differentiating cancerous and non-cancerous nuclei with a p-value of 1.02e-06. In addition, this study achieves an accuracy of 78 percent using a computational and machine learning-based pipeline for analyzing the morphological heterogeneity of nuclei and protein expression, enabling characterization of patient-specific tumor phenotypes. Unlike traditional morphology analysis pipelines limited to specific imaging platforms, this workflow enables cross-cohort and cross-cancer comparison, capturing tumor-specific phenotypic deviations at a single-cell resolution. The resulting phenotypic profiles could inform prognosis, treatment, and monitoring of therapeutic response.

**Summary:** As cancers become more aggressive and require more energy, typically uniform and organized cells begin to develop abnormal features to support their heightened needs. Studies have found that the prevalence of abnormal features is directly associated with the speed at which the tumor grows, but also the body’s ability to fight back. This study aims to streamline the analysis of these irregular features across all cancer types, providing a clearer picture of how nuclei distinguish stages of cancer and aid in rapidly clinically assessing at-risk or affected patients. Using the nuclear abnormality score developed, this study was able to identify sub-populations of highly irregular cancer cells, and successfully separate them with a p-value of 1.222e-21. By comparing the expression of cancer proteins with these irregularities, we can begin to develop insights that can be used across all imaging techniques to understand the cancer’s inner workings and learn to predict relapse before it even occurs.

## 1 Introduction

The introduction of chemotherapeutics by Goodman and Gilman’s in 1956 marked the beginning of modern oncology. Since then, strategies have become increasingly sophisticated, yet cancer relapse remains a defining limitation. In Multiple Myeloma (MM), for instance, the presence of a semi-solid polyclonal cancer population leads to a greater risk of remission over more homogenous malignancies such as breast cancer. As a result, MM’s 10-year overall survival is just 31.6 percent for patients diagnosed 2010–2015. Complete, sustained remission plunges from 32 percent after first-line therapy to only 2 percent by the fifth line (Wajs, 2013). Even when targeted therapies succeed against one subpopulation, resistant strains can emerge and proliferate, worsening patient outcomes (Wajs, 2013). Histology analysis, or the practice of examining tissue samples for clinical insight, has long guided clinical response against relapse, especially in Breast Cancer and Multiple Myeloma, where subtypes of cancer cells display unique nuclear features (Fujino, 2018). Since 1855, researchers like Rudolph Virchow have used the distinct morphological traits present in cancer cells to identify tumor cells and adjust cancer treatment (Kampen, 2012). In Leukemia, specifically, Virchow was able to diagnose his first patient using histology in 1846, identifying and distinguishing based on the presence of strange, abnormal cell boundaries and nuclei in leukocytes (Kampen, 2012). In a modern context, histology can also allow clinicians to identify which “clone” of cancer cells caused the disease and how far the tumor has progressed into key tissues such as the bone marrow, a key prognostic indicator. However, despite centuries of innovation, these conventional tactics have been limited by subjectivity, variability between observers, and labor-intensive workflows that restrict scale and reproducibility. To address these limitations, modern molecular methods like single-cell RNA sequencing have been developed offering greater resolution through transcriptomic profiling. While powerful for characterizing cell identity, these methods often sacrifice spatial context, affordability, and morphological detail, essential factors for understanding tumor architecture, immune evasion, and clonal expansion. Furthermore, single-cell data is often full of noise and difficult to validate, as samples are sensitive to manipulation and RNA data often expresses highly complex signals. Imaging Mass Cytometry (IMC) and higher-dimensional cell imaging, on the other hand, combine multiplexed protein detection with high-dimensional spatial imaging, capturing structure and expression at a single-cell resolution. In this study, we develop a pipeline that integrates nuclear morphological profiling with protein expression to investigate the structural hallmarks of aggressive and potentially resistant breast cancer clones, allowing for ‘at-a-glance’ analysis of a patient’s entire tumor, and functioning regardless of the type or stage of solid cancer. By correlating nuclear irregularity with cancer progression, we identify a link between structural phenotype and malignant behavior. In addition, we identify a population that we denote cancer progression “drivers” composed of distinct large, highly abnormal cells that alter the non-cancerous cells around them. Beyond therapeutic inefficiencies, a major issue in cancer tissue analysis is the potential for misdiagnosis with an average rate of 8.4 percent for all diseases, and a problem that severely injures over 800,000 lives in the United States each year (Newman-Toker, 2024). In a modern clinical context, cancers specifically are diagnosed through a sequence of techniques ranging from initial physical exams to imaging techniques, and ultimately through a biopsy (Pulumati, 2023). Despite the rigor of current diagnostic methods, misdiagnosis remains a major issue, with 1 in 7 breast cancers misdiagnosed by doctors (Ryser, 2022). Whether a false positive or a relapse, misdiagnosis by a clinician often means serious injury and in worse cases unnecessary loss of a family member. In order to drive down rates of misdiagnosis, tools that integrate tried and tested strategy with machine intelligence must be developed to work in concert with clinicians achieving human adaptability and machine accuracy all in one pass. Taking inspiration from Virchows discoveries and the field of histology this study aims to analyze the most important morphological marker of mutation in the cell: the nuclei. By quantifying nuclear irregularity we hope to eliminate subjectivity in cancer histology and catch hidden trends within cancers that could save more lives without forcing clinicians to invest more time. In summary, This project aims to define general “key” cancer identification metrics and create an easily implemented pipeline to understand the morphological differences between cancer and healthy cell lines across all solid cancers. We hypothesize that a small set of nuclear morphological features can robustly distinguish malignant from benign cells in all cellular imaging types and solid tumors, setting the path for future clinical analysis.

### 1.1 Study Goals

- Create a pipeline for cell abnormality analysis based on images of cell nuclei.
- Find distinctive nuclear morphological features in cancer cells that can help differentiate malignant cells regardless of image or cancer type.
- Streamline comprehensive histological analysis for clinicians using high-dimensional imaging data and key metrics
- Train a machine learning model to identify cancer cells in situ while also identifying non-cancerous cells of note

## 2 Methods

### 2.1 IMC Data Preparation

First, according to the Danenberg study, breast cancer samples were taken from patients, and segmented into sheets of 2-6 micrometers (Chang, 2017). Tissue sections were paraffin-embedded to maintain structural integrity during processing. Metal-conjugated antibodies targeting proteins such as Histone and Cytokeratin (CK) were applied according to established IMC protocols. These antibodies facilitate elemental labeling detectable by laser ablation in the Hyperion CyTOF system (Danenberg, 2022). The plate of affixed cells were then placed in a Hyperion CyTOF machine, where the concentration of specific proteins was estimated using mass spectroscopy and the amount of specific metals that react and ablate following exposure to a laser. Protein concentrations were quantified based on metal intensity, overlaid on eachother, and saved as multi-channel TIFF image stacks (Danenberg, 2022).

### 2.2 Single Mask Analysis

Using the *imread* function, we made this full-stack TIFF file visible and preprocessed it to achieve consistent contrast and visibility. We then generated a cellular and nuclear mask using a pretrained segmentation model. We then extracted marker channel labels, and the preprocessed the full stack channel corresponding to a cancer protein and displayed it as an RGB mask on top of the cellular mask. We then calculated the per-cell mean RGB value, and we put the expression of each marker into a tabular file. Using Python, we calculated the solidity, aspect ratio, circularity, and other abnormality metrics for each cell and aggregated them into a single “abnormality score” that allows for simplified analysis and separation.

**Table 1:**
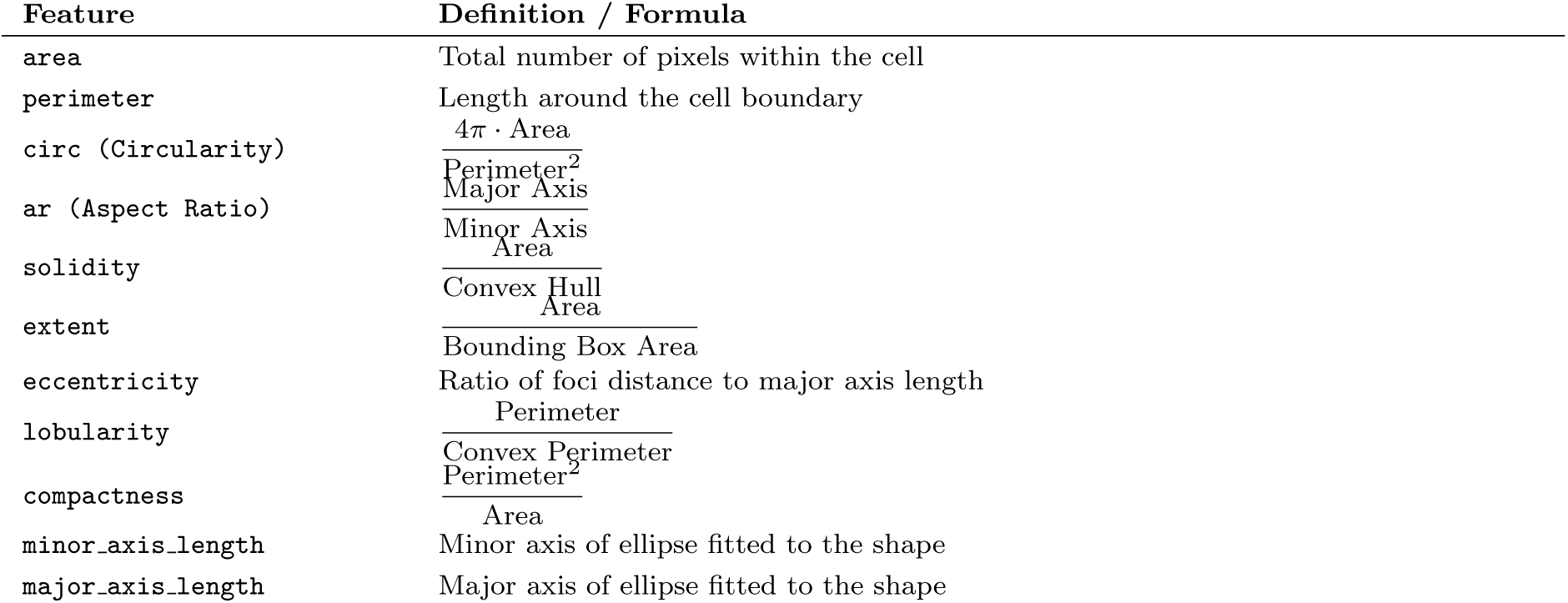
Definitions and formulas of shape-based features used in model.

Using the package *region props* from *skimage.measure*, each nucleus was then identified, with the area and perimeter being calculated using the *region.area* and *region.perimeter* functions. Finally, a *dataframe* was made that stores all marker and morphological signals, allowing for data analysis and corresponding plots to be generated.

To ensure metric accuracy and effectiveness, we created visual plots displaying sample cells to allow for quality control and combat potential missegmentation. In addition, we overlaid per-cell heatmaps displaying irregularities on the original mask to provide additional quality control. These heatmaps were then compared with cancer masks and markers to ensure that the abnormality provided a direct correlation to cell state.

**Figure 1:**
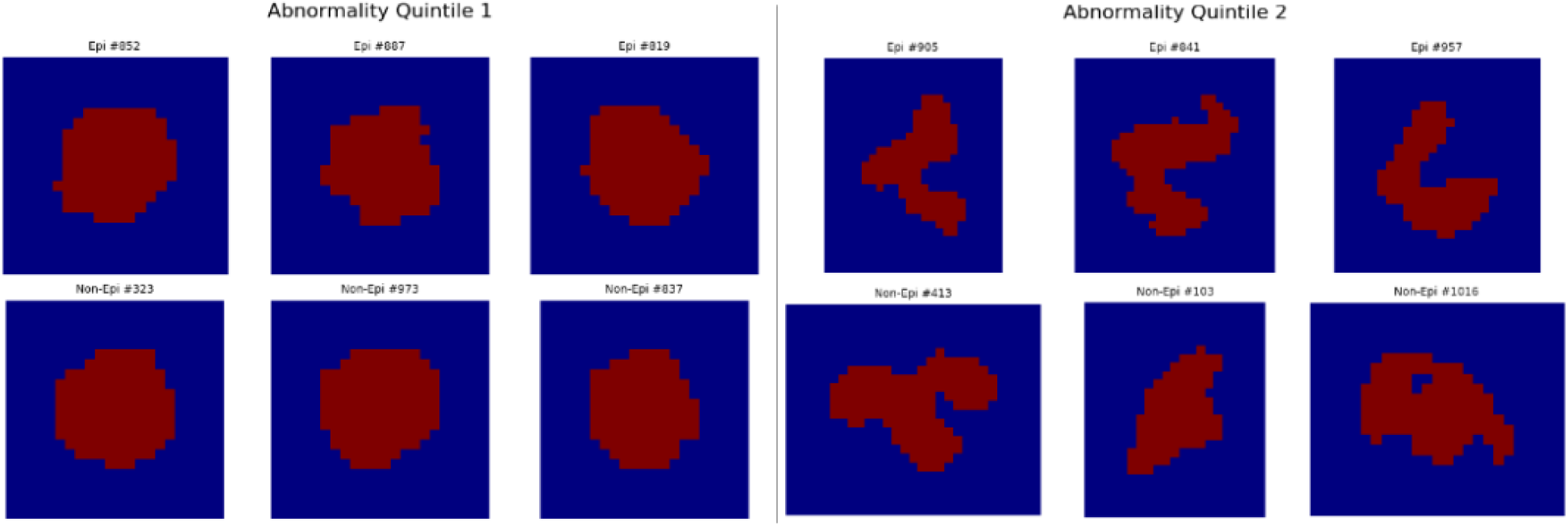
Visualization of the distinct difference between highly Abnormal (Quintile 2) and Normal Cells (Quintile 1) using their nuclear mask and grouped based on Abnormality score

These formulas allowed for circularity and abnormality scores to be generated based on the masks created by the cellular segmentation model. Morphological scores were overlaid onto the original segmentation masks to visually confirm the spatial distribution of nuclear irregularities.

**Figure 2:**
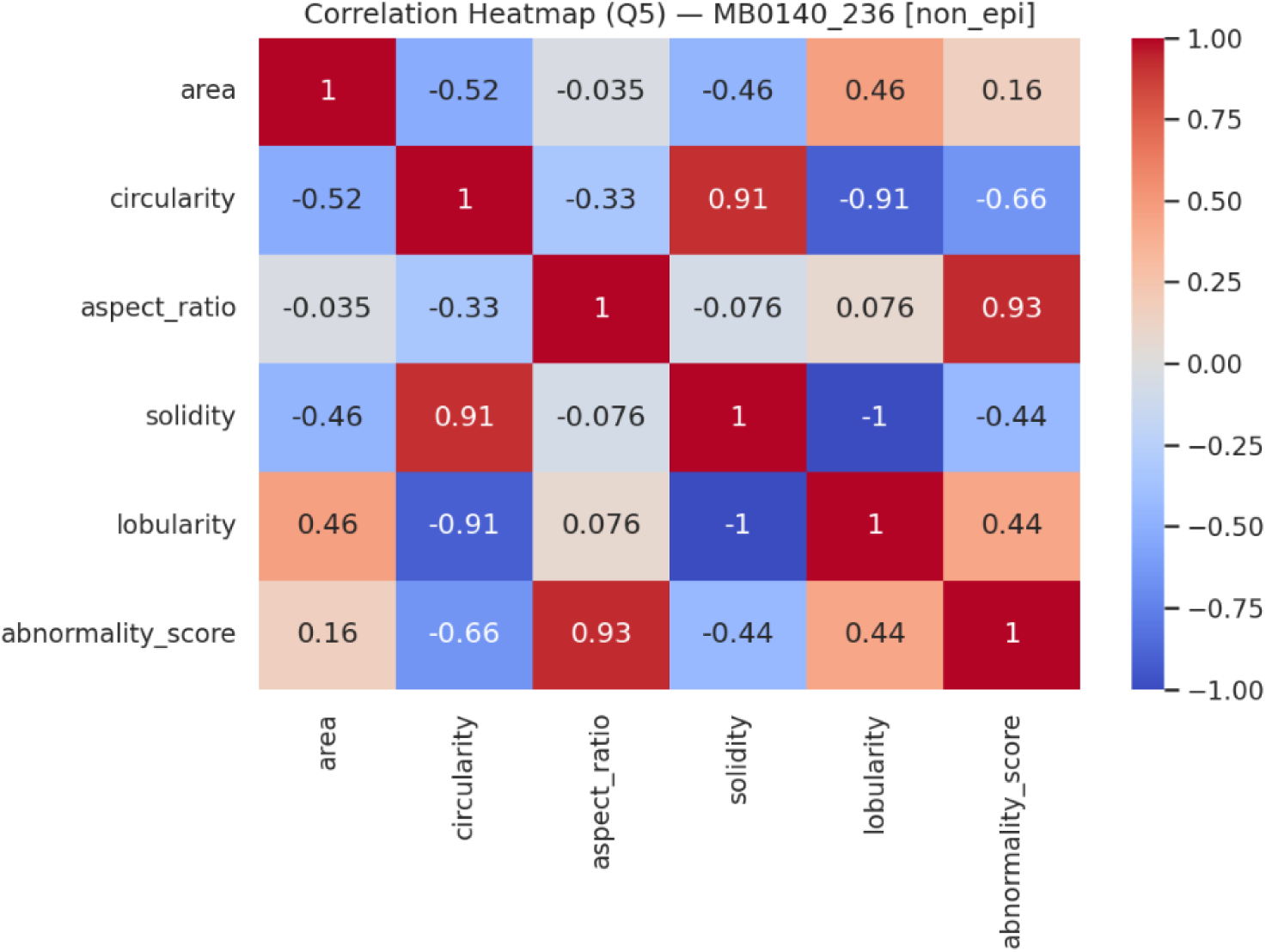
Simplified Correlation Plot Cell Metrics represent relationships between several key metrics and their impact on the final abnormality score

### 2.3 Minimum Cell Size analysis

While Imaging Mass Cytometry deals with subcellular micrometer-level resolution, the average diameter of an animal cell is only 30 micrometers. Any outliers below a certain area value suffer from inaccuracies in abnormality values due to the limited resolution for perimeter/area calculations. To mitigate resolution-induced measurement errors, nuclei with an area below 20 pixels were excluded from the final data frame.

### 2.4 Viable Sample Selection

We screened approximately 400 IMC samples from the 2022 Danenberg study. We selected the 10 with the most balanced cancer and non-cancer ratio and pre-processed for further analysis, while also “removing” sub-threshold nuclei. This included removing cells that were over 300 pixels squared in area, and under 20 pixels squared in area, as we visually identified these cells to be populations (only 0.5 percent of all sampled) of missegmented cells.

**Figure 3:**
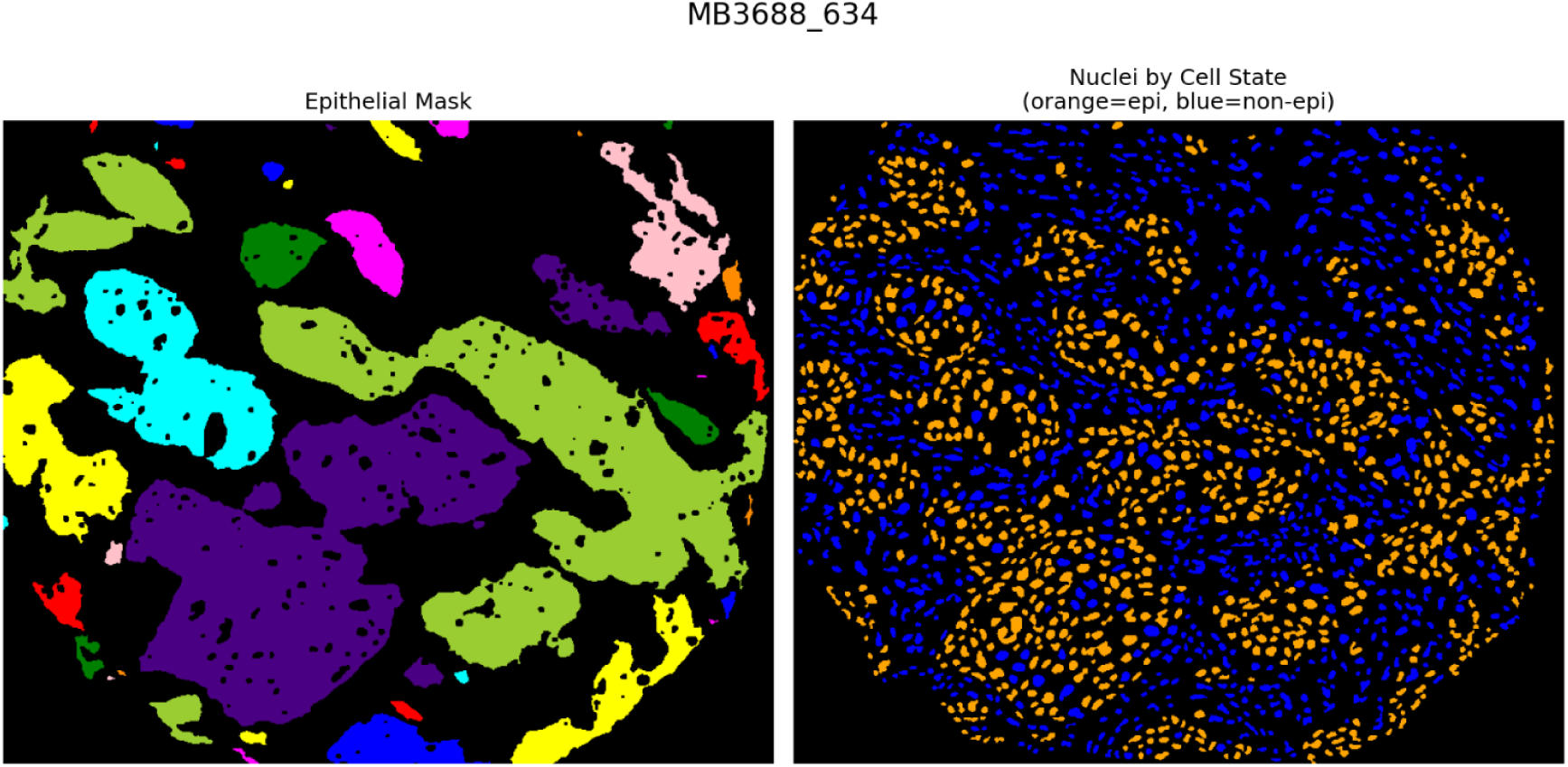
Epithelial mask overlaid on nuclei displaying how Viable samples are selected

Group Mask Analysis Using Google Drive, we connected the three main folders to Google Colaboratory: Nuclear Masks, Epithelial Masks, Full Stack Masks, and Cellular Masks. Using a “for” loop and the unique “MB” code for each sample, we paired the masks to their corresponding samples (Danenberg, 2022). WE then processed the masks separately, with the nuclear mask used to identify morphological shifts across the different cell types identified by the markers (taken from the full mask) plotted on the cellular mask (Danenberg 2022). The epithelial mask, on the other hand, was overlaid on top of the nuclei, with all nuclei under the mask being labelled as cancer. By combining these masks, we were able to match files and morphological features across all cell types allowing for analysis and comparison.

### 2.5 Multiple Mask Analysis

While individual mask analysis is important to identify themes, more generalized data was also needed, so we generated a histogram for circularity and lobularity using Matplotlib across the 10 viable nuclear masks available at (Danenberg, 2022). In addition, as a control for the relationship of different metrics, a pair plot was generated, and all significant markers were independent of area, and markers of abnormality correlated correctly.

### 2.6 Cancer Cell Identification

First, we used the markers that corresponded with cancer cells, such as *HER2* in breast cancer. We overlaid an RGB mask on top of the nuclear mask and compared the abnormality of cells in the HER2 “cloud” to those that were outside. Although marker-based identification achieved promising results, signal spillover, where highly expressing cells increase the signal of surrounding cells, common with surface markers like *HER2*, limited specificity prompted the use of spatial epithelial masks for more reliable classification and imaging type-agnostic analysis. Using the mask created by the Danenberg study we were able to define the cancer cells by comparing the overlap of the mask to the centroids of their nucleus. This allowed for a binary comparison where non-cancer and cancer cell lines could be compared both on their nuclear morphological features, but also their expression of certain proteins/markers. An added benefit was the ability to generalize the nuclear analysis to any type of cellular imaging, as it relied on simple images of nuclei rather than hard-coding the analysis of specific proteins and genes.

### 2.7 Quintile Analysis

To identify themes and populations of interest across all masks available in the Danenberg study, we generated quintiles (bins) to separate high-abnormality and low-abnormality cells. We classified each cell across the total population into percentiles with 20 percentile intervals representing “bins”. As represented in Figure 4, all cells were classified into quintiles regardless of cell state, allowing for ratios and cell-state based differences to be calculated across bins.

**Figure 4:**
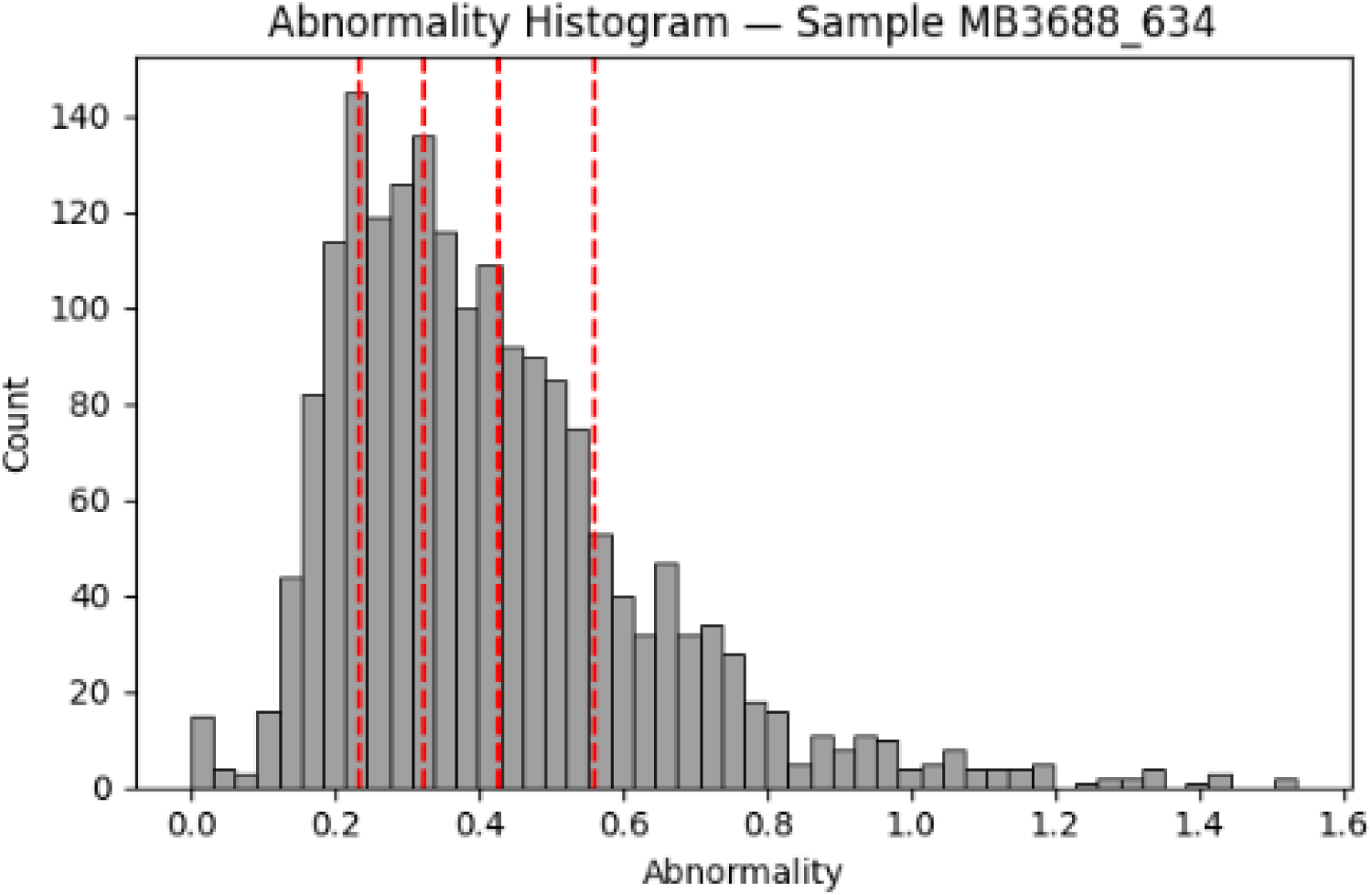
Histogram of abnormality values across quintiles demonstrates presence of clear abnormal cell “tail”, and distinct populations across a viable sample

To separate the most from the least abnormal cells, we generated bins based on abnormality score and area and compared to see the phenotypic differences in nuclear morphology. To ensure homogeneity, features such as Area and distribution of nuclear features were compared to ensure that no confounding variables could influence the expression of nuclear markers.

### 2.8 Random Forest and XGBoost Analysis

Although not all epithelial cells are cancerous, the majority of cells within the epithelial masks are expected to be cancerous in this context. To ensure that epithelial cells that we analyzed reflected cancer change rather than just differences in cell type, the epithelial masks were compared against cancer marker images, as show in in Figures 21–23. Only samples in which 80-90 percent of epithelial cells were cancerous were included. Having done this quality control, all cells with nuclei under the predefined “cancer” mask were marked as cancerous, and a Random Forest and XGBoost model were trained on these labels to identify potentially cancerous cells with abnormal nuclei. The signals between classes were then compared and used to identify important metrics for diagnosing cancer.

### 2.9 Data

All data and formulas were then exported to a GitHub repository for usage by other researchers in analyzing cancers in their labs https://github.com/hamcoderfran/MM-IMC-Tool. All data was written in a Jupyter notebook combined with Google Drive to allow for streamlined analysis based on file name (*NuclearMask* and *EpithelialMask*).

## 3 Results

Layering a cancer marker RGB mask (Figure 5)and quantifying the mean expression of that RGB value across a nucleus and setting gates for “abnormal expression,” we were able to identify cancer cells using markers with high accuracy (78 percent) in a preliminary trial. However, when repeated and combined with morphological analyses, this marker and nuclear shape analysis achieved an even more accurate separation of cells of over 85 percent compared to the epithelial masks.

**Figure 5:**
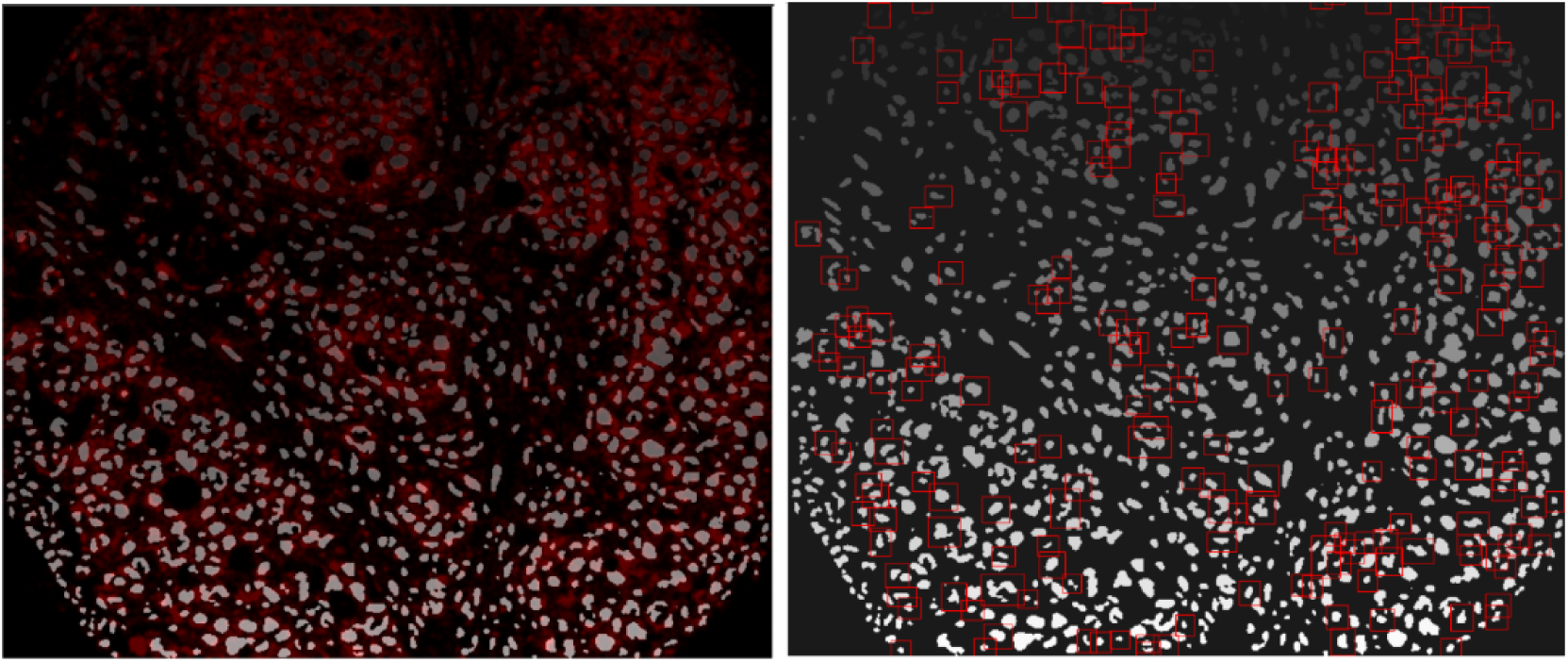
Visual of how marker expression is used in gating cancer versus non-cancer cells through marker expression

However, as seen in the Figures 6 and 7 below, while HER2 is a viable metric for detecting cancer versus non-cancer, complex segmentation models are necessary as the simple gates set in this study were not generalizable. There was no significant change in HER2 expression between classes except for in circularity. When applied to a scatter plot, certain values were extremely highly expressed while others were down-regulated. In addition, the potential for extremely high-expressing cells and errors in segmentation made the usage of markers too variable, and ultimately the correlation was non-significant. To resolve this issue, we used the existing Gaussian mixture model (GMM)-generated epithelial masks from Danenberg (2022), allowing the workflow to be adjusted to any segmented nuclear data.

**Figure 6:**
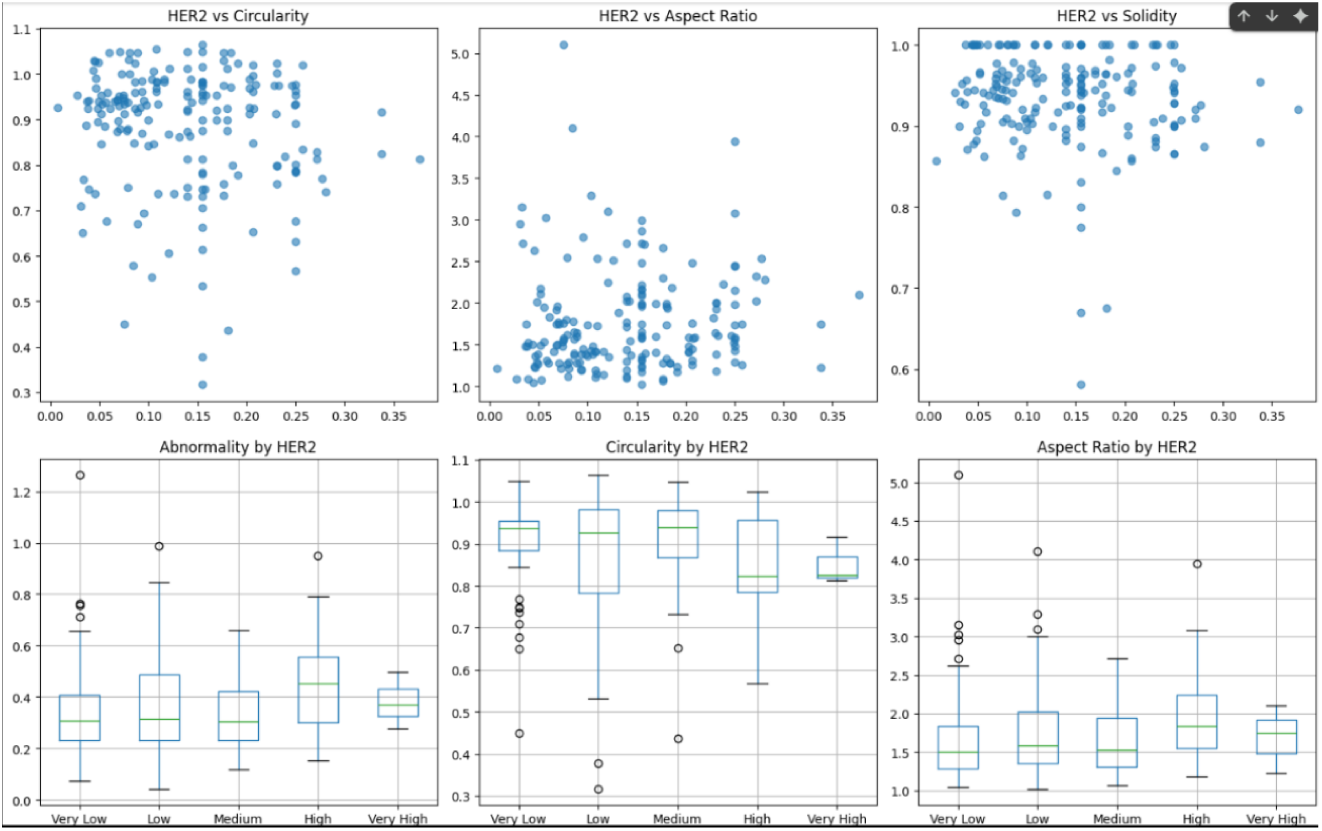
Lack of coherent trend in HER2 expression across abnormality scores except for circularity

**Figure 7:**
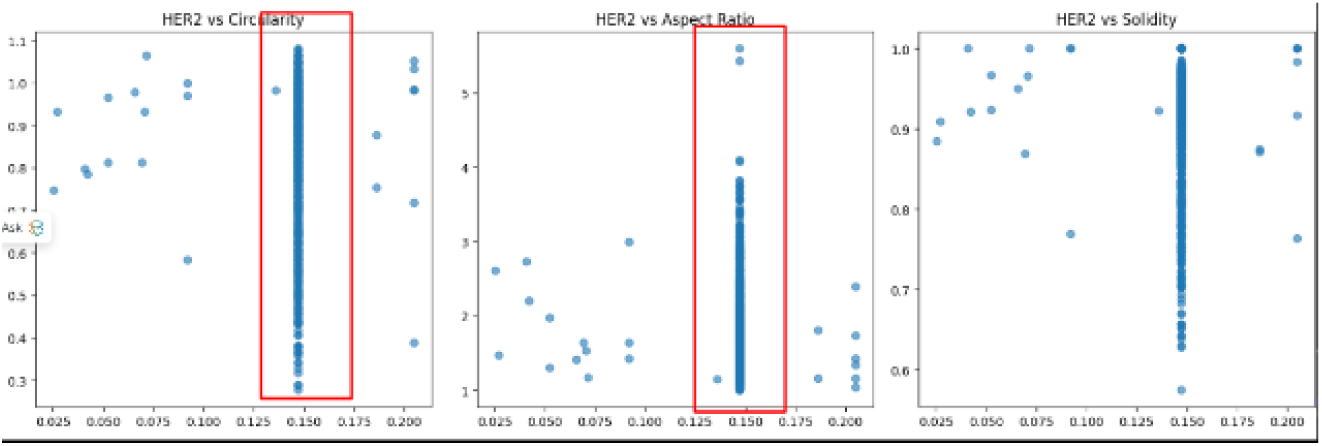
HER2 signal spillover compromises marker-based classification fidelity across samples and limits ultimate classification accuracy

On the other hand, Epithelial masks were able to consistently generate a statistically significant mean/median abnormality difference between healthy epithelial and non-epithelial cells (Figure 8). Using a Mann-Whitney U test, the masks achieve a statistically significant difference in 10 out of the 10 viable masks, and consistently were below a p-value of 1e-05. To ensure the epithelial mask accuracy, nuclei separated by cancer state were visually compared against cells expressing high amounts of BC marker, and there was no significant difference between the two.

**Figure 8:**
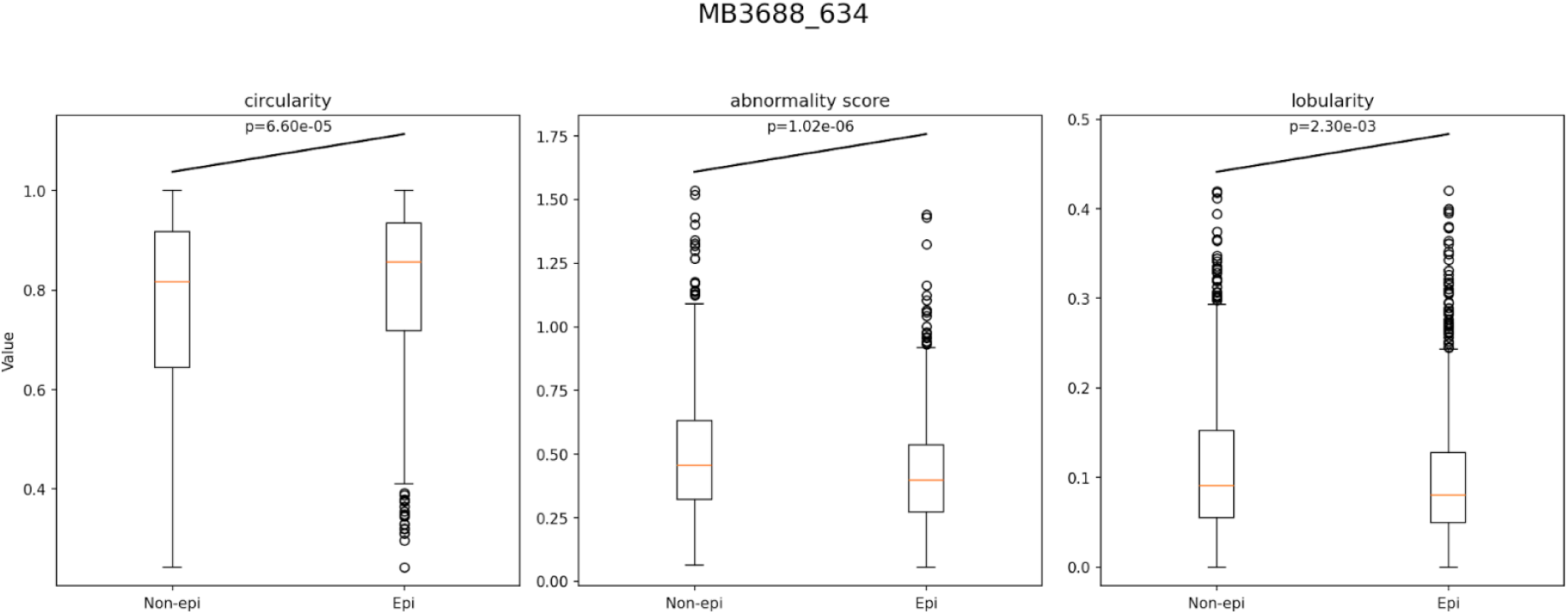
Boxplot of Cancer (Epi) and Health (Non-Epi) displaying the statistically significant increase in abnormality

Using epithelial mask analysis (Figure 8), we found a statistically significant difference in cancer cells’ abnormality across all samples with p-values consistently below 0.001, such as 2.73e-04 in MB3412, 5.78e-07 in MB5718, and 1.02e-06 in MB3688. This demonstrates a class wide differences based on cell state, created by abnormal cancer mutation and folding of the nuclear membrane.

In addition, we identified the key population driving this difference in the form of a high area high abnormality population indicated in Figure 9 by purple Q5 cells, which represent the cells that are most abnormal and compose the tail seen in Figure 4 and 20. When comparing this to the box plots of epithelial and non-epithelial ratio and Figure 20/17, this population is mostly composed of cancer cells, indicating that both the metric and workflow can consistently identify the most malignant cells within cancer, and separate them. This is further supported by the statistically significant epi/non-epi ratio (Figure 13, which shows that q5 represents mainly cancerous cells, identifiable through morphology analysis.

Stratifying cells into quintiles based on abnormality scores revealed that the top 20 percent (Q5) were significantly enriched for cancer cells, exhibiting distinct morphological signatures and higher nuclear deviation metrics. As seen in Figure 10 and 14, there is a substantial difference between cell types in terms of abnormality and morphological scores, indicating the importance of the cancer state in the final nuclear morphology.

**Figure 9:**
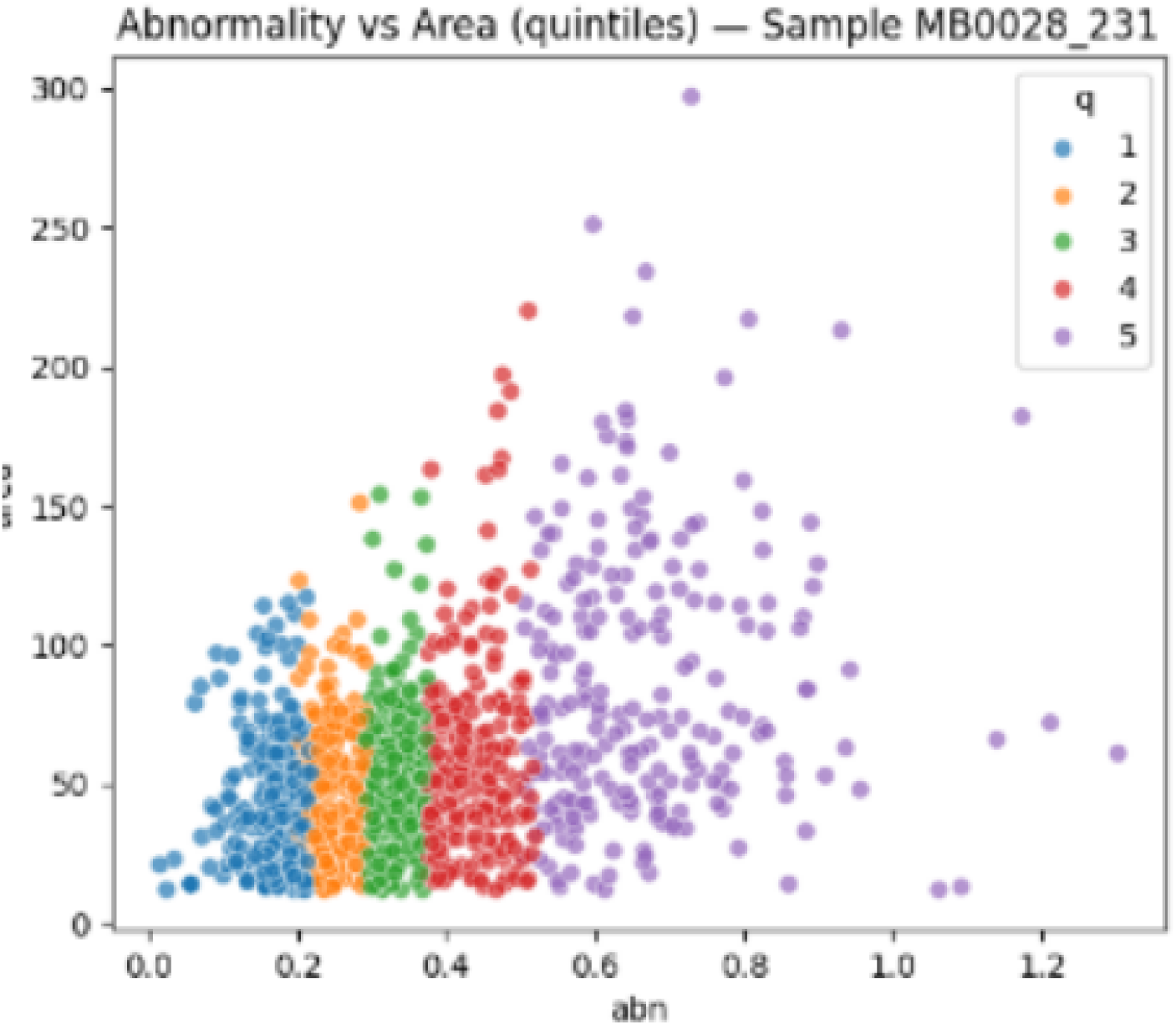
Quintiles representing different cell populations across abnormality score

**Figure 10:**
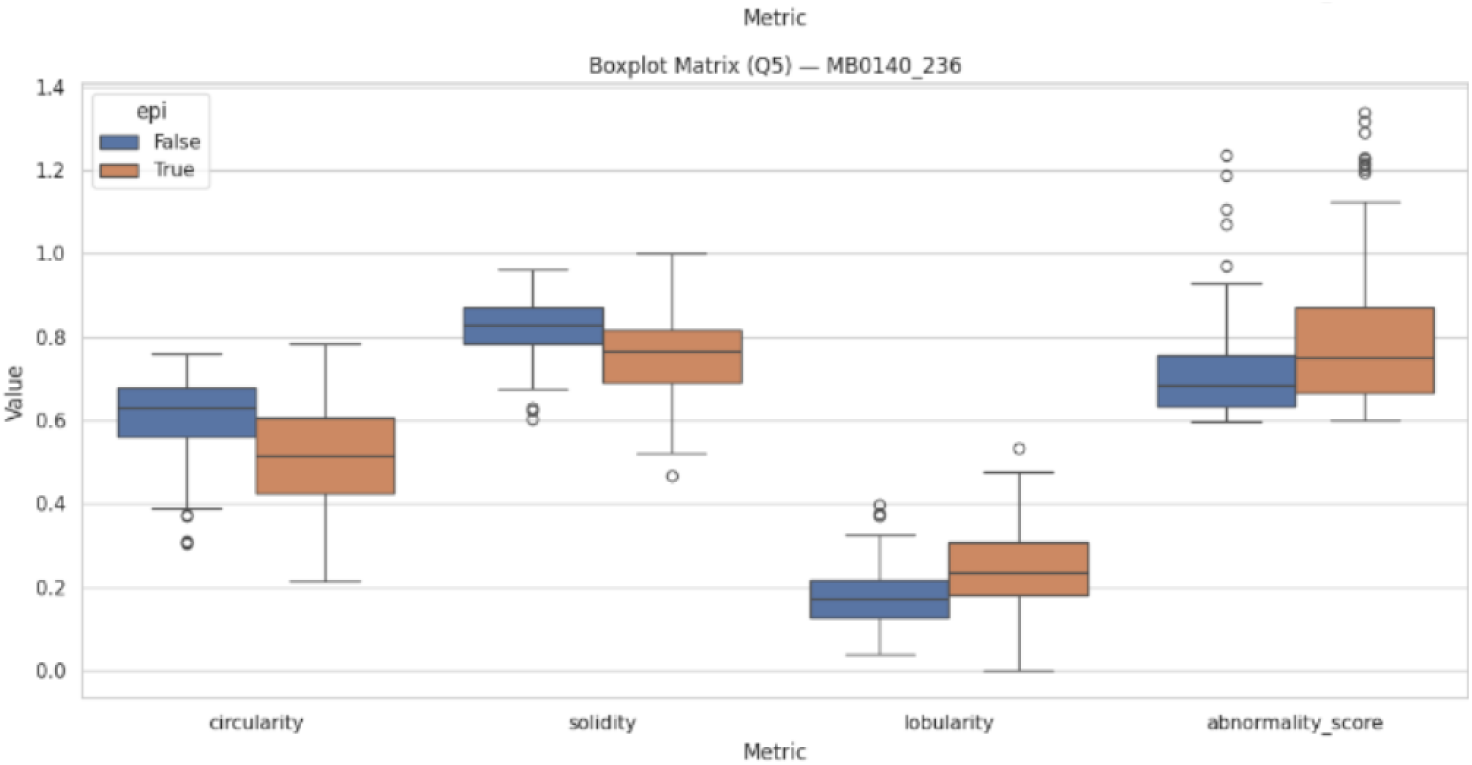
Difference in metrics across cell types in the top 20 percent for abnormality

This cell-type-based difference based on quintile is made more apparent when cells are separated by class and abnormality and plotted against area. A distinct pattern emerges, with a similar area across all cells in quintiles 1 and 2 (20 and 40 percentile), and a significantly higher area for cancerous cells in q5. In addition, the slope of abnormality appears different across cancer and non-cancer, influenced by the high “abnormality” cancer cells in the upper quartiles. This is further shown in Figure 11 and 15 where there are distinctly separated epithelial cells with high area and abnormality driving up the slope of the line of best fit and demonstrating a different progression towards abnormality than is typical in breast cell lines. This could indicate the presence of TME-altered cell progression causing even non-cancerous epithelial cells to develop abnormal features according to tumor cues.

**Figure 11:**
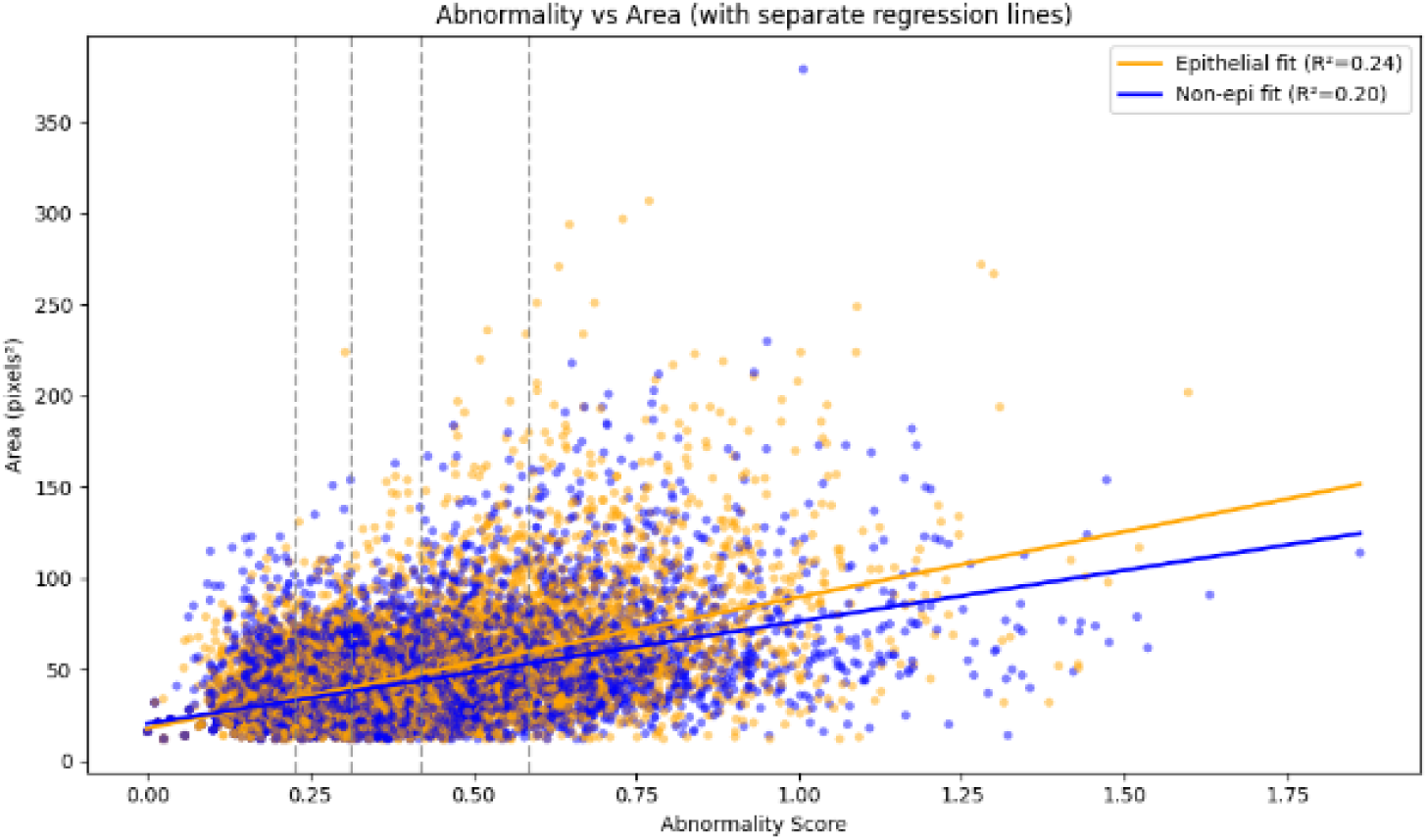
Plot of distinct abnormality quintiles across samples

To confirm this larger presence, the ratio between cancerous and non-cancerous cells was calculated, and as seen below in Figure 13, the quantity of epithelial cells substantially increased as the bins selected for increasingly abnormal cells. This demonstrates the metric’s ability to select for cancerous cells at a higher level as they progress further and further highlighting the ability for the tool to point out pockets of highly progressed cancers versus less aggressive Q1-Q3 cancer cells. Across all bins in Figure 13, the median ratio between cancerous and non-cancerous cells increased from 0.75 in the lowest abnormality quintile to 0.9 in Q4 and 1.25 in Q5. This demonstrates how the abnormality metric is both increasingly able to differentiate abnormal cells as they mutate and progress, but also that abnormality can distinctly pick up the presence of cancer in the form of epithelial breast cancer cells. The scale of this difference is further shown in figure 14, where the p-value displays a statistically significant difference between cell classes even in Q1 (p = 1.05e-16), Q3 (p=2.57e-37), and Q5(p=1.22e-21). When comparing by Area, a key feature of hypertrophied breast cancer cells emerges where advanced cancer cells begin to expand their nuclei, and this shift is reflected by the extremely high cancer/non-cancer ratio in Figure 12 and 19 where just in Q3, Q4, and Q5 there is a higher quantity of epithelial cells as area increases (Narasimha, 2013). This adds further evidence to the notion that these metrics are able to increasingly select for abnormal cells that express key cancer and malignant development markers as they become more pronounced.

**Figure 12:**
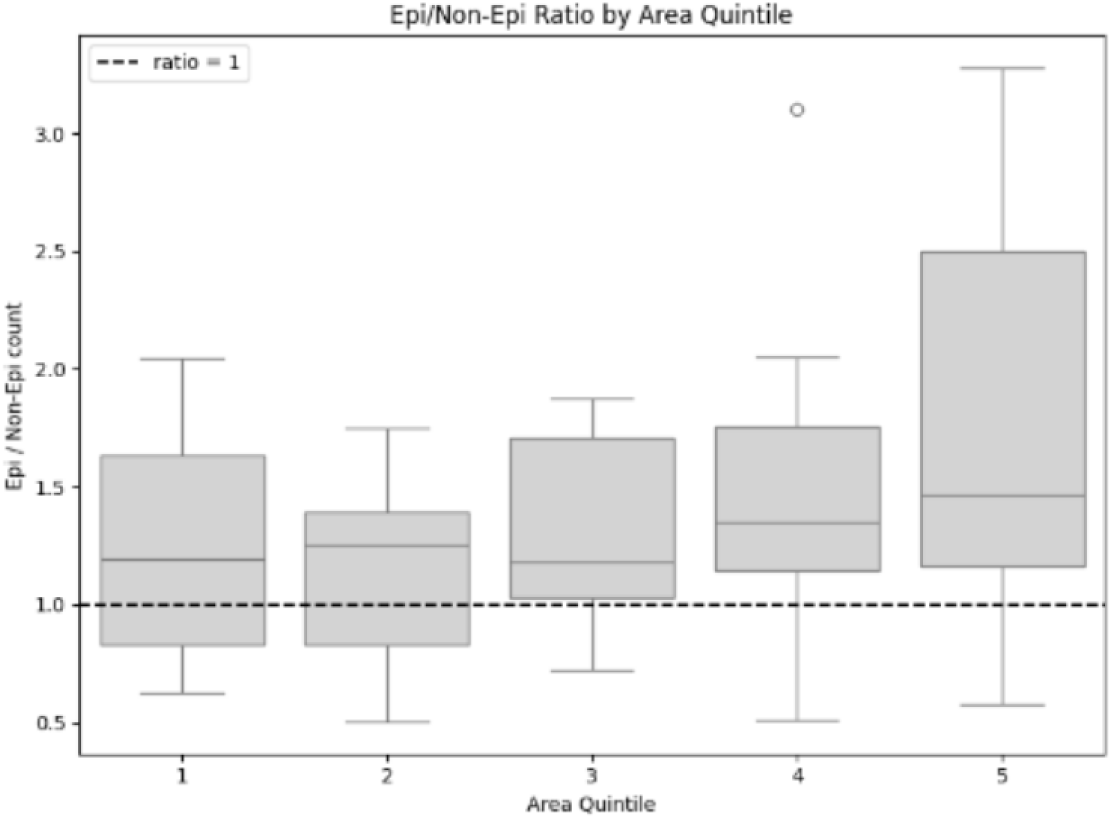
Cancer to Non-cancer ratios in quintiles based on area displaying significant

**Figure 13:**
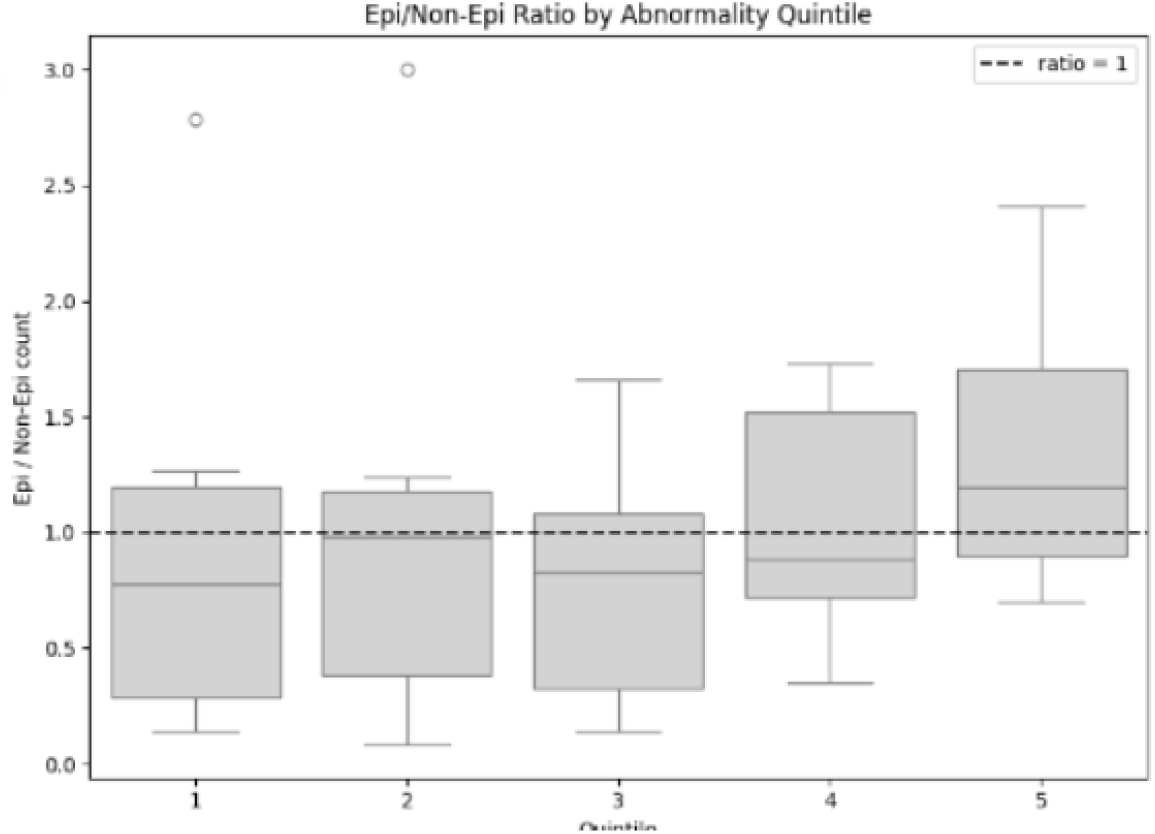
Cancer to Non-cancer ratios in quintiles based on abnormality

Figure 14 further demonstrates this divide with a p-value of 1.22e-21 between the epithelial and non-epithelial (cancerous and non-cancerous) cells in the most abnormal cells in Q5, and 1.06e-62 in Q4. In addition, Figure 14 highlights the capacity of the abnormality score to separate cancerous cells at a higher and higher level as they become more abnormal, while also keeping a statistically significant difference of 1.05e-16 and 1.74 e-33 at lower abnormality quintiles. This adds further evidence to the capacity of these metrics to separate cells based on cell state, even if they aren’t the most abnormal in the sample. Figure 14 and 11 adds more context to this difference, demonstrating how the difference in abnormality between cell types increases as they become increasingly abnormal, and indicating these metrics’ ability to pick out unhealthy cells. Further support is given in Figure 15, which displays a dimensionality reduction plot where the distinct tails of highly abnormal epithelial cells, are distinct from non-epithelial cells representing the different paths of progression and influence of cancer on the expression of certain morphological metrics.

**Figure 14:**
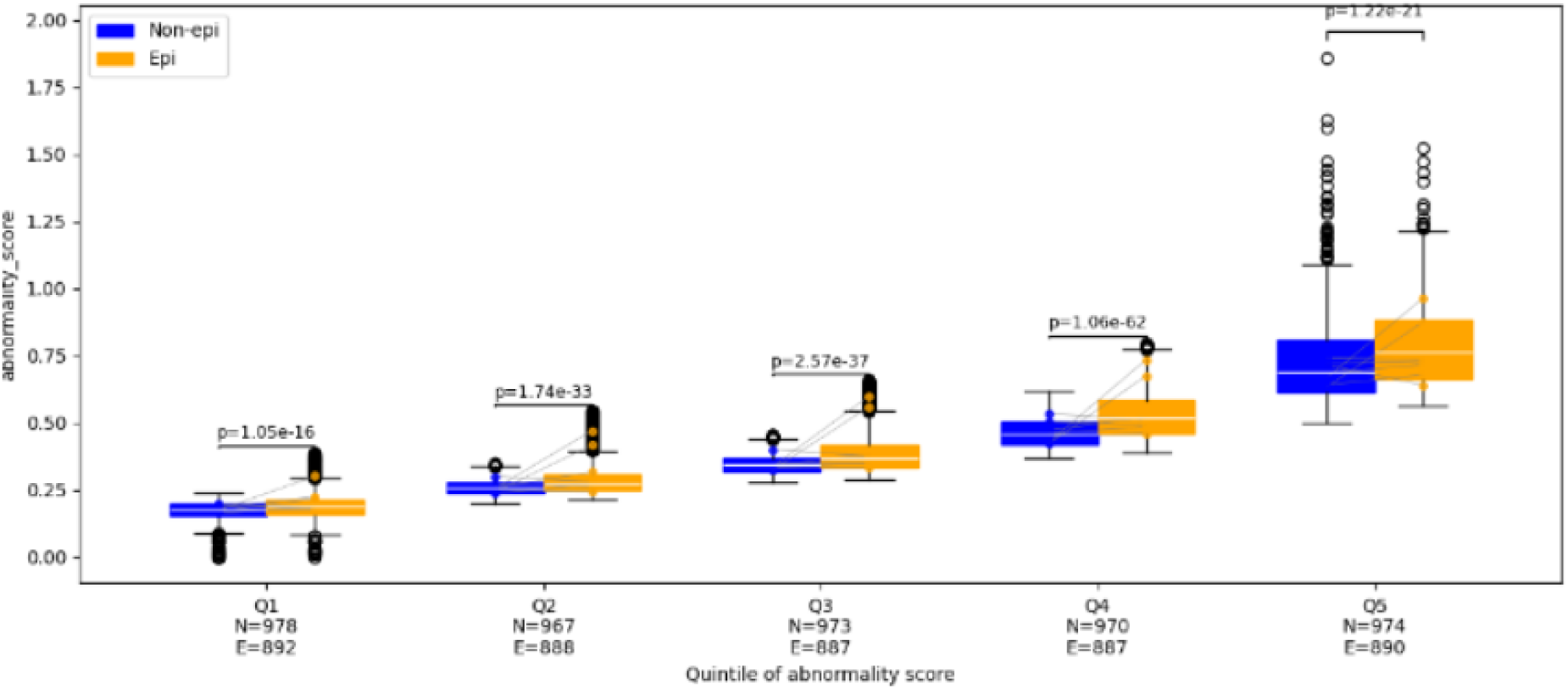
Abnormality Boxplots across all samples and Quintiles

**Figure 15:**
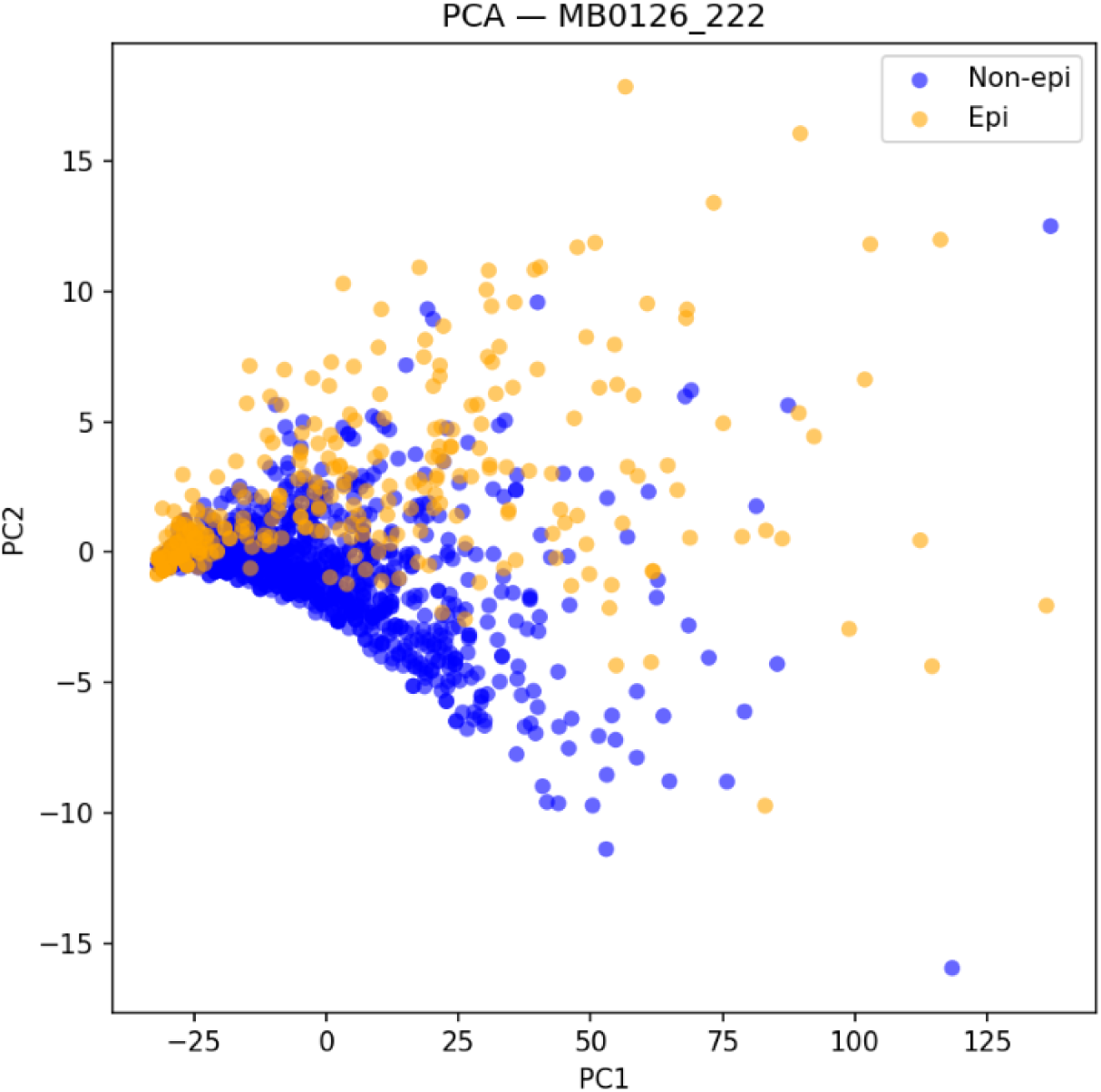
Cell type wide differences between epithelial and non-epithelial cells demonstrating different progressions and distinct phe-notypes

In addition to a supervised and human-sourced analysis through an epithelial mask, an unsupervised machine learning approach was also included. Random Forest and XGBoost models were trained on nuclear features to classify cancerous versus healthy cells. To ensure generalizability, the model was only trained on 10 samples, which were deemed viable, and tested on a separate of 2 images. An XGBoost model trained on extent, aspect ratio, solidity, and other morphological metrics was able to achieve an accuracy of 0.74 and AUC of 0.71. A Random Forest model with similar training data was able to achieve an accuracy of 0.72 and an AUC of 0.65. To ensure that the metrics were not misrepresenting the tools ability to seperate cancer and non-cancer images were generated of predictions (Figure 16), to ensure that certain segmenting and staining errors weren’t driving down accuracy due to procedural errors.

**Figure 16:**
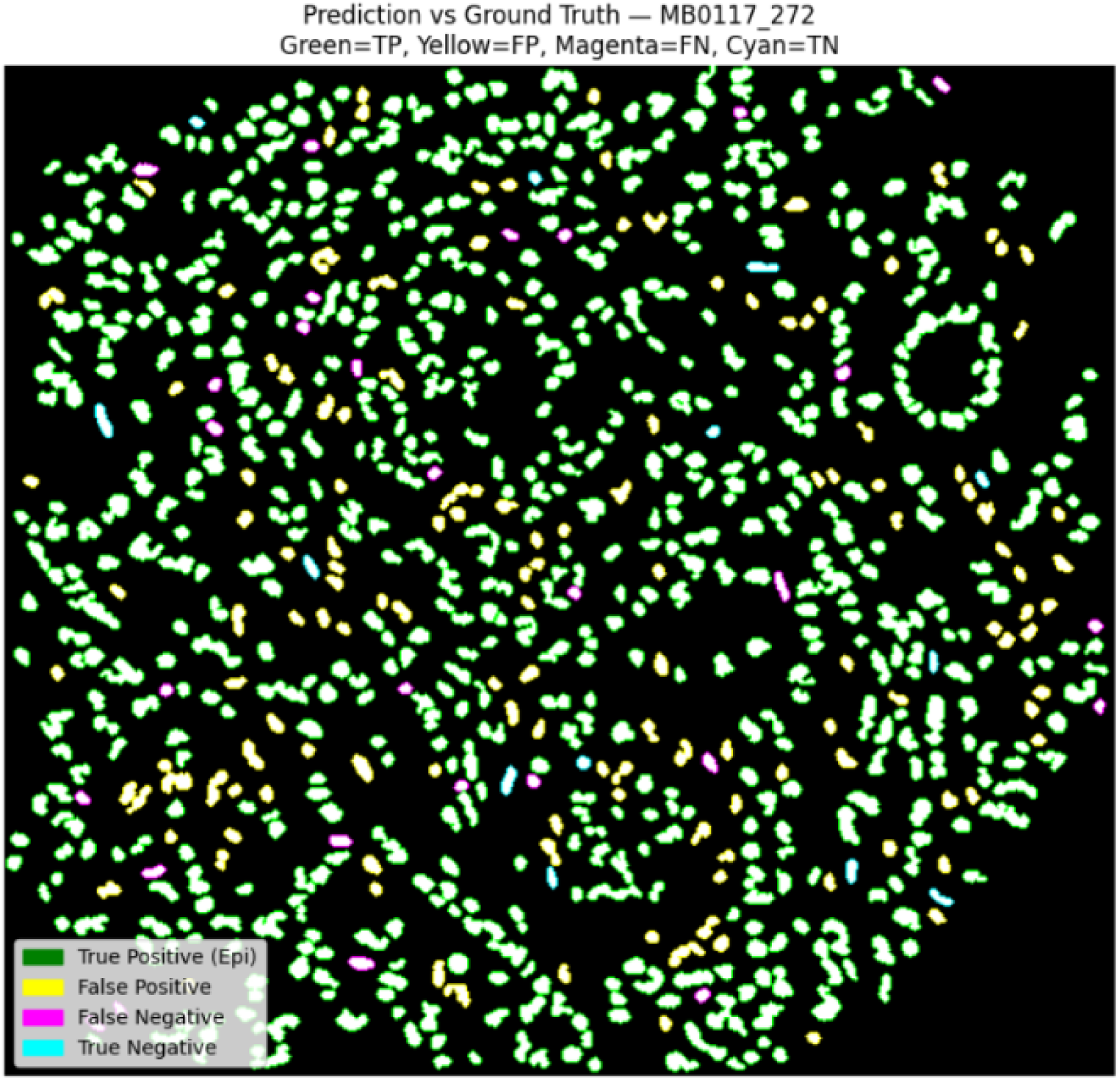
Image of XGB model accuracy across a nuclear mask displayed using colors

## 4 Discussion

A central goal of this study was to ensure that image-derived abnormality metrics were able to differentiate between normal non-epithelial cell characteristics and those of directly cancerous or cancer influenced epithelial cells. As both the *HER2* and *ER* marker-based mask and epithelial mask display a considerable correlation with cells with much higher abnormality scores (Figure **??**, this indicates that the metric can pick up on abnormalities generated through cancer progression rather than simply the inherent qualities of being epithelial. The presence of a highly abnormal 5th quintile cancer cell population (Figure 17) indicates that there is a significant malignant population in breast cancer that can be separated purely by selecting for the top 20 percent in terms of abnormality. When combined with other metrics, this can then be generalized further to other quintiles allowing us to separate cancer cells across all abnormality bins with a high degree of accuracy In figure 18, this is made evident with the 5th quintile being primarily made of epithelial cells, which are more abnormal on average, and follow a separate trend of progression far steeper than the healthy non-epithelial cells. This adds evidence to the idea that cells comprising the TME (epithelial), could potentially follow a cancer-altered differentiation trajectory, where normal cell organization and nuclear development are altered by surrounding oncogenic signals and TME-guided cues. Epithelial cells as a whole may adopt aberrant phenotypes, with certain aggressive populations driving the overall expression of abnormal size and shape of nuclei, ultimately shaping a pro-tumor niche.

**Figure 17:**
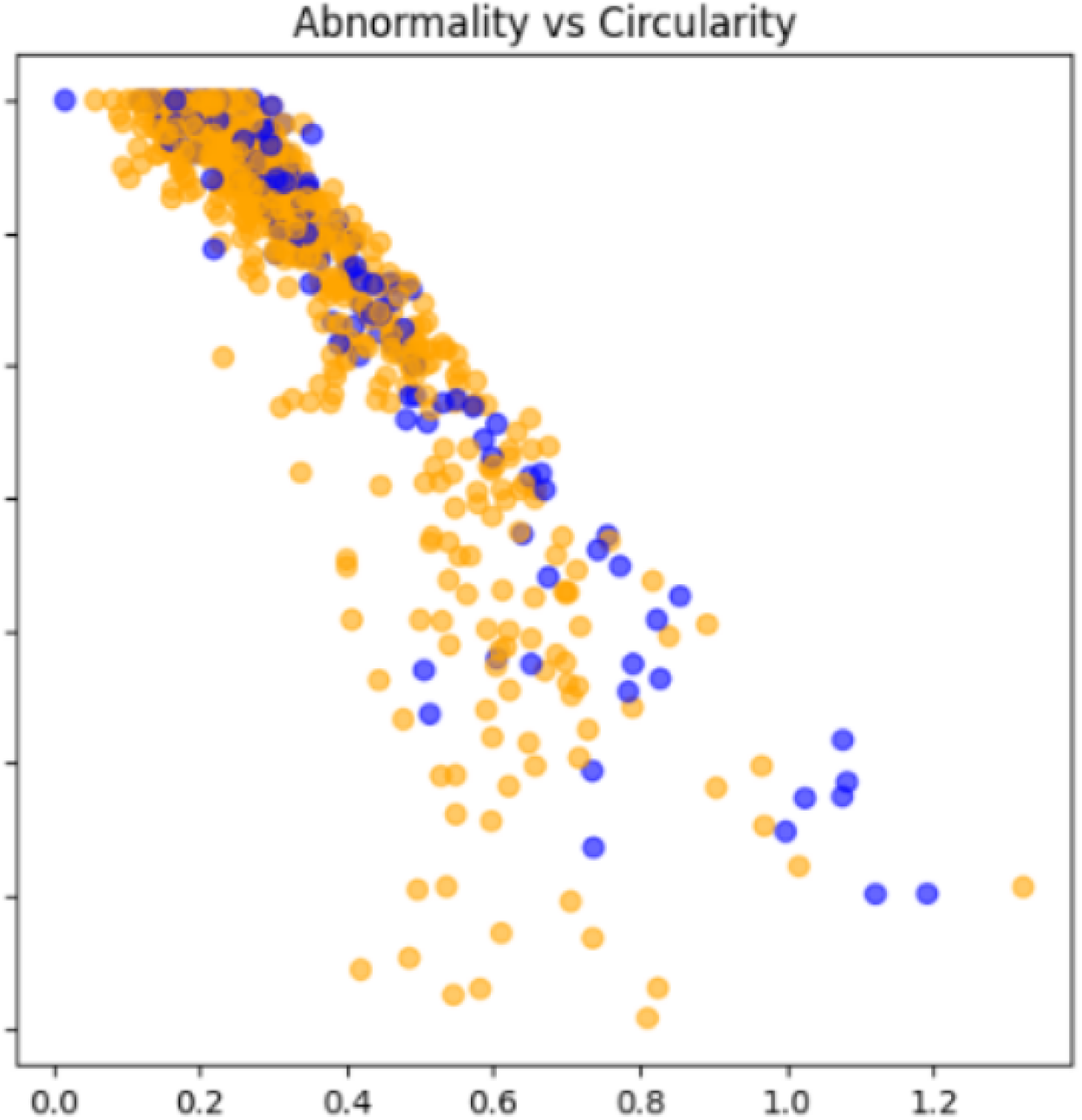
Cells plotted by abnormality and circularity

Although epithelial masks formed the backbone for classification in this study, integrating multimodal features could further enhance diagnostic precision and clinical applicability.

Ultimately, the Random Forest model was able to achieve an AUC of 0.65 and an accuracy of 85 percent across epithelial classes. The XGBoost model performed better with an AUC of 0.71, and a similar accuracy of *<* 0.75.

Notably, minor axis length emerged as the most predictive feature in the XGB model. This aligns with prior studies showing that elongated nuclei are indicative of DNA replication stress, weakened chromatin tethering, and altered lamina composition, all of which contribute to tumorigenicity (Singh, 2022). When visualized, cancer cells demonstrate statistically significant (*p <* 0.01) elongation compared to non-cancerous counterparts, adding further evidence that the minor axis may be a pan-cancer signal of malignancy. There was a substantial difference between populations with an average minor axis of 6.74 pixels in epithelial cells and 5.59 pixels in non-epithelial cells (Figure 18).

**Figure 18:**
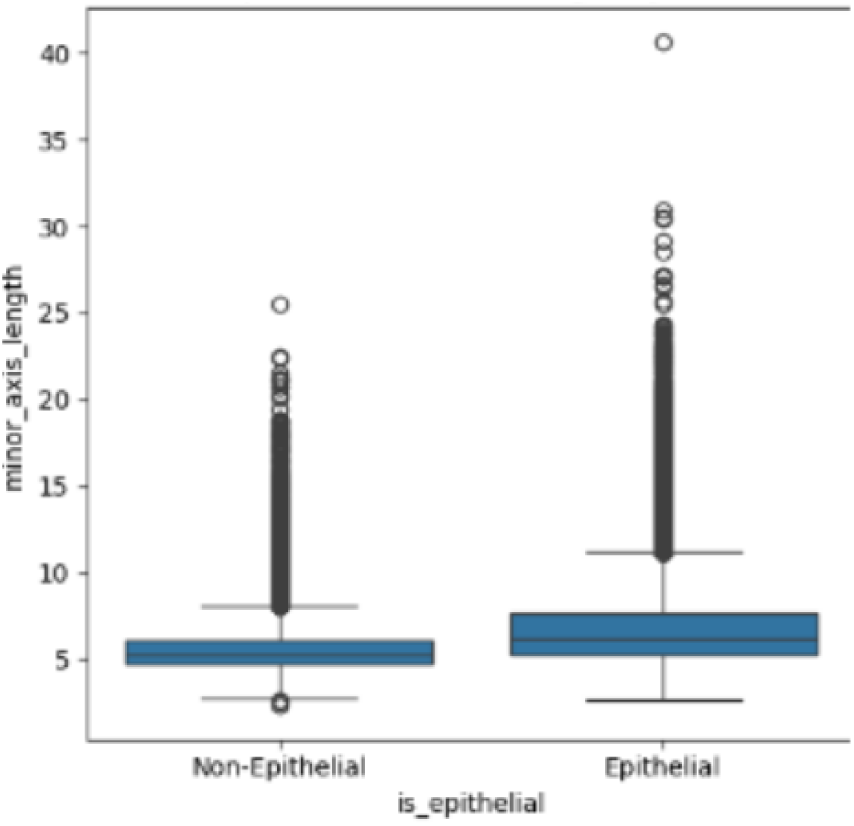
Minor Axis length across Cancerous and Non-Cancerous cells

Further validation was provided through area-based stratification. Nuclei exceeding 100 pixels squared consistently showed greater overall abnormality as seen in figure 13 and 20, reinforcing the widely held idea that larger structurally irregular nuclei represent a signature of aggressive cancer cells. These measurements may reflect hypertrophy, the dysregulation of nuclear import/export dynamics, or abnormal ploidy features, which are all implicated in chemotherapy resistance and genetic instability (Denais, 2014). Such pleomorphic metrics could be used in detecting tumor prognosis in aggressive cancers like melanomas, with the NIH finding that greater nuclear hypertrophy and abnormality in the invasive band reflects a more aggressive tumor behavior and a higher potential for metastasis (Bian, 2024).

**Figure 19:**
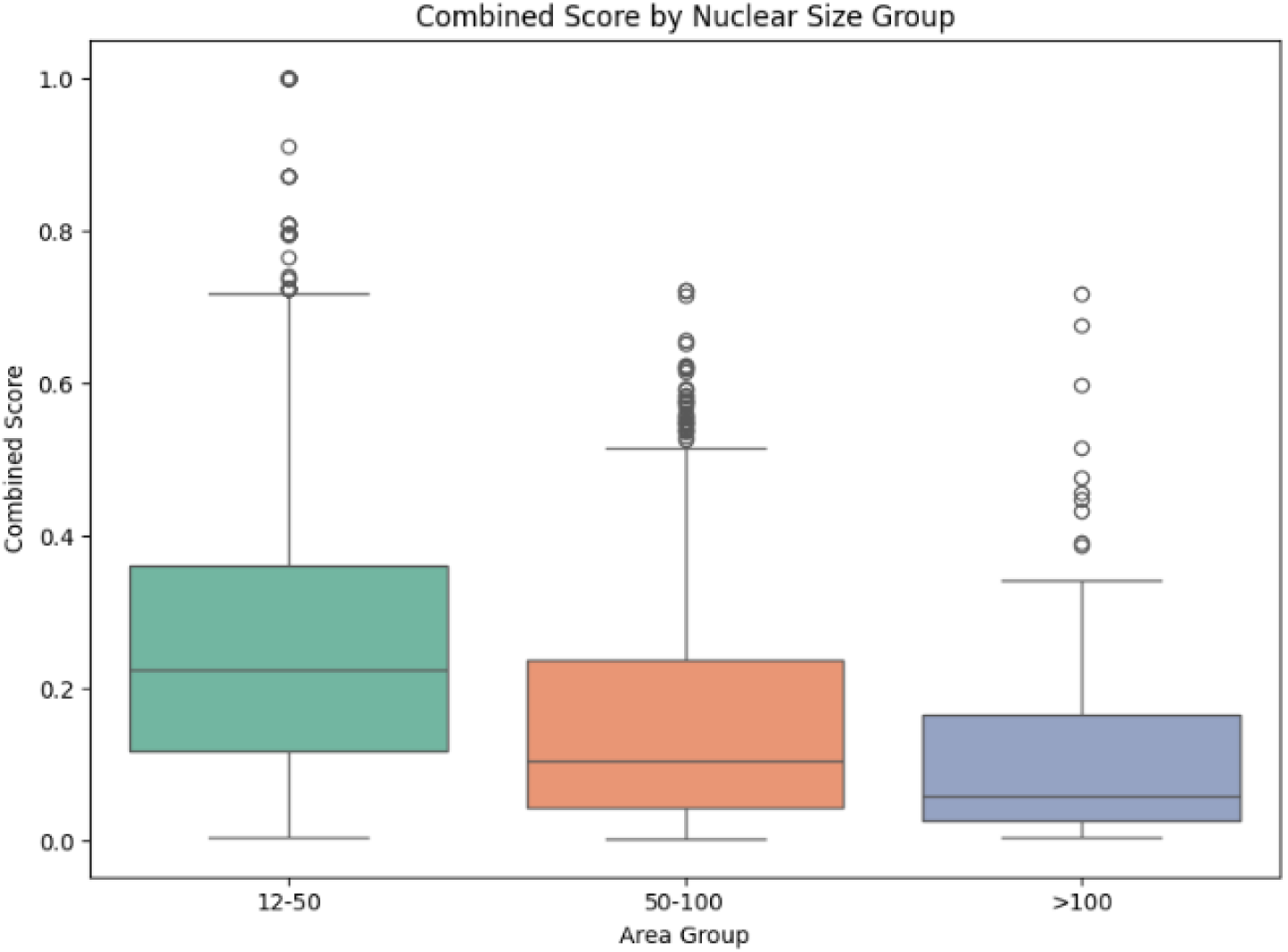
Abnormality as affected by area

Aspect ratio and solidity provide additional classification power. Cancerous epithelial cells are more elongated and less compact forming a morphometric separation that is visually separable from healthy cells, as we see in Figure 20 and 1. Across all these metrics, a clear tail of highly aggressive cancer cells emerges across the PCA and histogram plots (PCA: Figure 15, Histogram 4), which express high levels of abnormality even compared to other epithelial cells. Crucially, this population points to the potential for morphology to not only describe cells, but also predict their capacity to divide and worsen patient outcomes. In the future, this population of highly abnormal and large cells can be used to infer metastatic potential, estimate transcriptomic shift, and reflect intracellular tension states across diseases.

**Figure 20:**
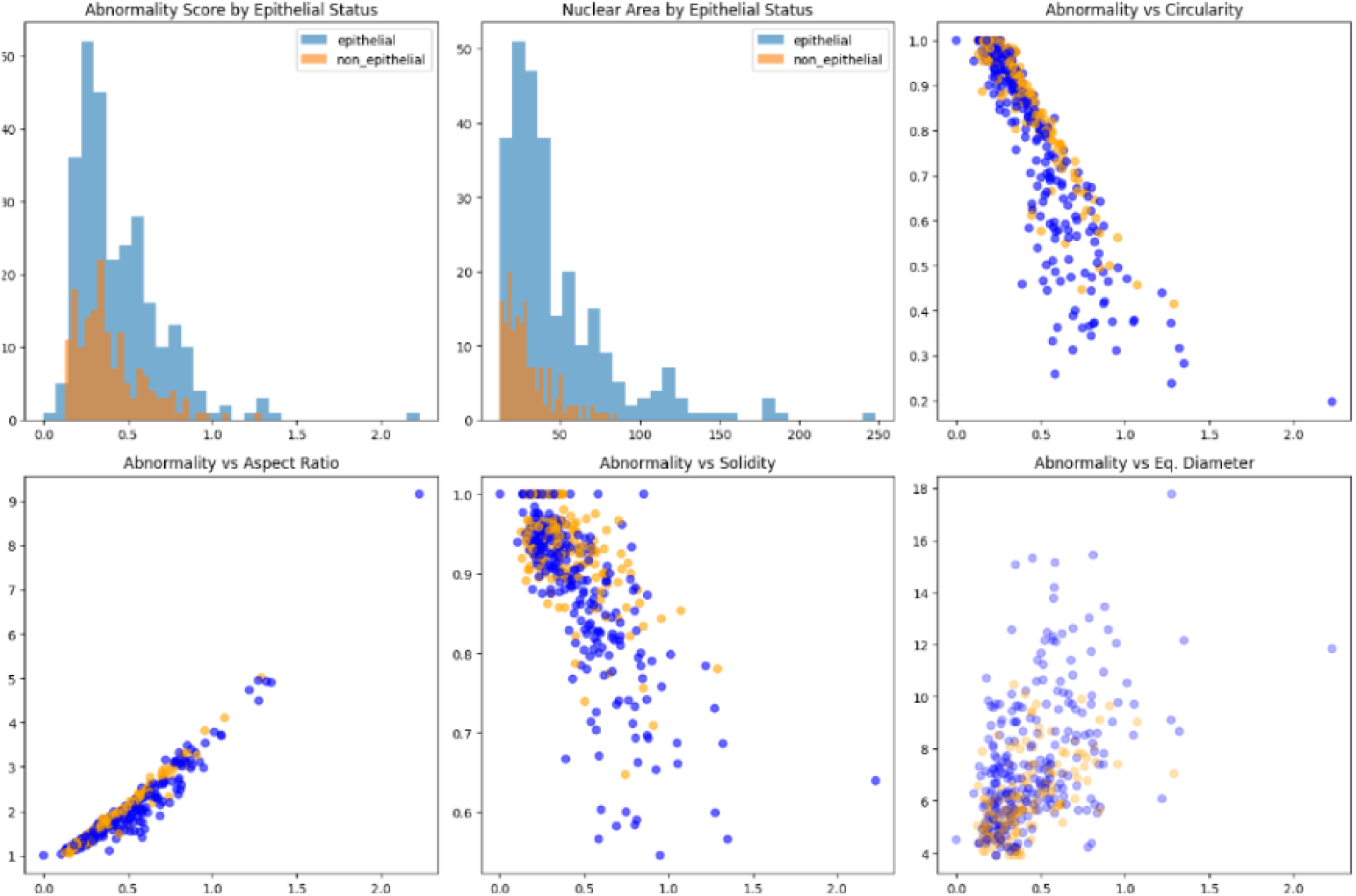
Seperating cancer and non-cancer cells based on abnormality metrics

To visualize hypertrophy and the importance of minor axis, a sample that has distinctly separated cancer non-cancer populations was plotted and compared against HER2. As seen in Figures 23 and 22 nuclei with high expression of cancer markers (HER2) also have extremely enlarged nuclei representing how larger nuclei may serve as a visible, quantifiable hallmark of malignancy. This reinforces how structural deformation and nuclear size serve as a proxy for underlying dysfunction.

**Figure 21:**
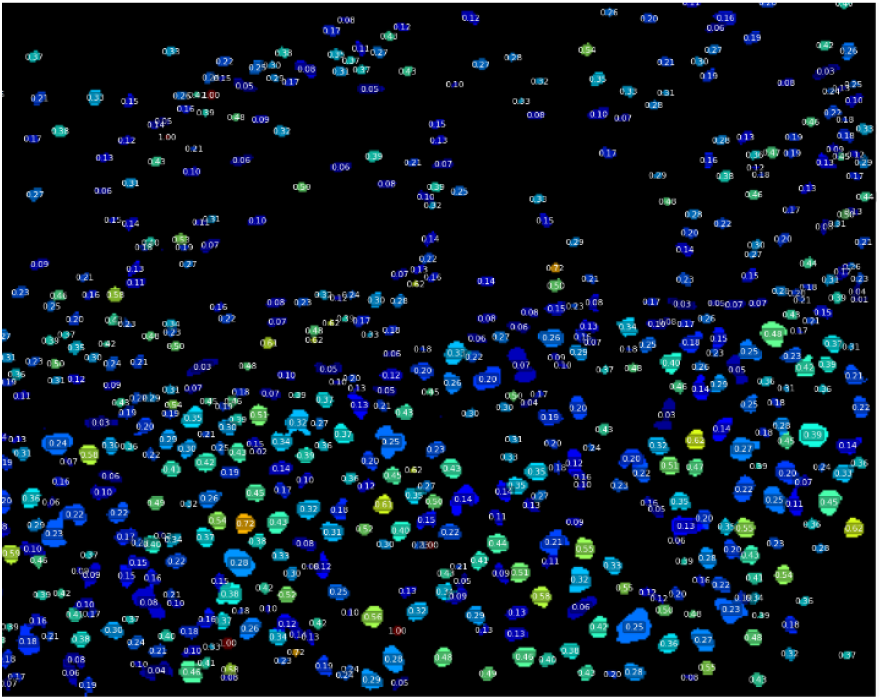
Clearly separated cancer and non-cancer samples show distinct differences in size and circularity, demonstrating hypertrophy in cancer cells.

**Figure 22:**
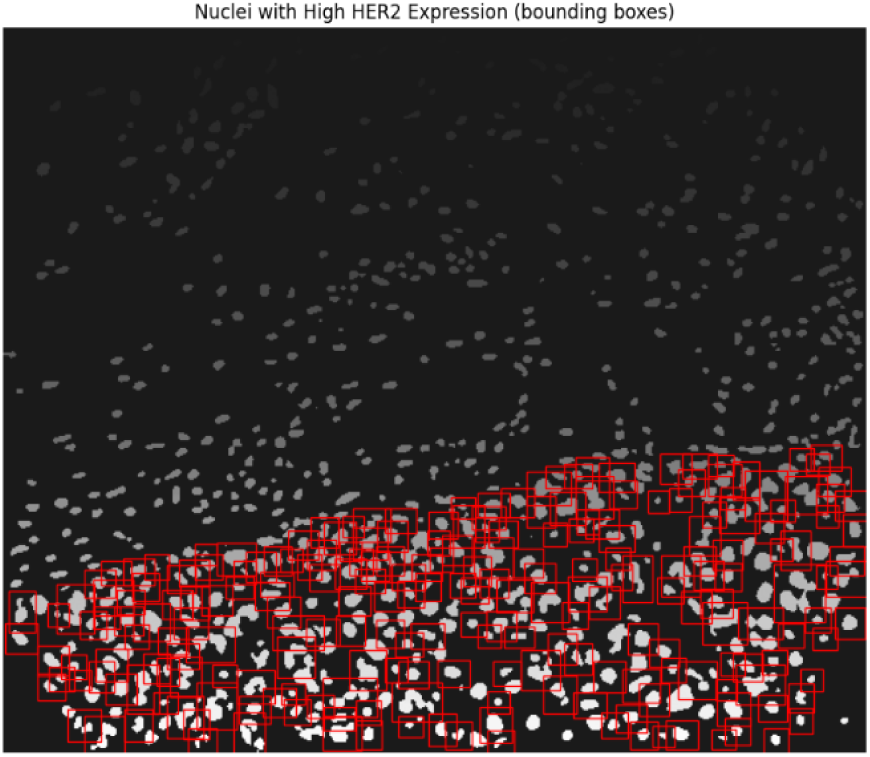
Nuclei with high cancer marker expression display increased nuclear area following cancer progression.

**Figure 23:**
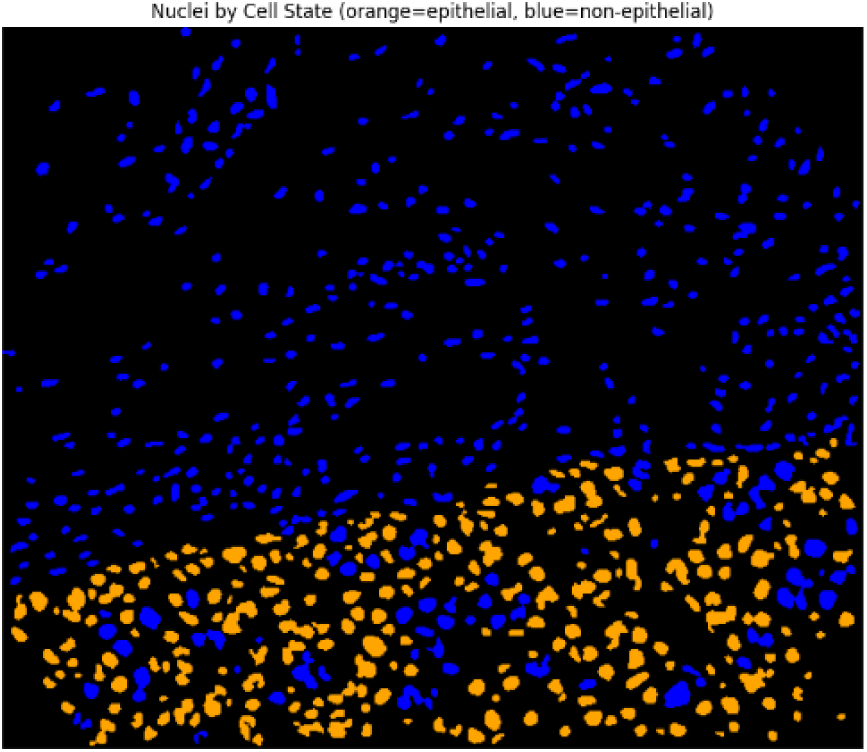
Epithelial mask overlay demonstrates masks ability to detect cancer cells as identified by a cancer marker

Interestingly, a common theme emerged in 8 out of the 10 viable samples focused on in this study: “pockets” of highly abnormal epithelial cells grouped together surrounded by abnormal non-cancerous cells. This mirrors the emerging understanding that a TME can be comprised of multiple clonal outgrowths, expansions of genetically or epigenetically distinct subpopulations, with polyclonality potentially being a marker of regions of the tumor with more resistance to therapy or co-evolution with vascular and immune cells. From a histological standpoint identifying pockets of abnormality surrounding cancer cells could help infer tumor aggressiveness and mapping cellular architecture in situ can reveal clinically significant zones within the tumor, informing diagnosis, risk of recurrence, and sites for targeted intervention.

### 4.1 Considerations

A main concern with the nuclei morphological analysis was the potential for total area of the nucleus to be a confounding variable due to the different sizes of nuclei present. To understand the effect of this variable the correlation between nuclei and abnormality score was calculated using correlation heatmaps, scatter plots, and histograms. No statistically significant correlation was found.

**Figure 24:**
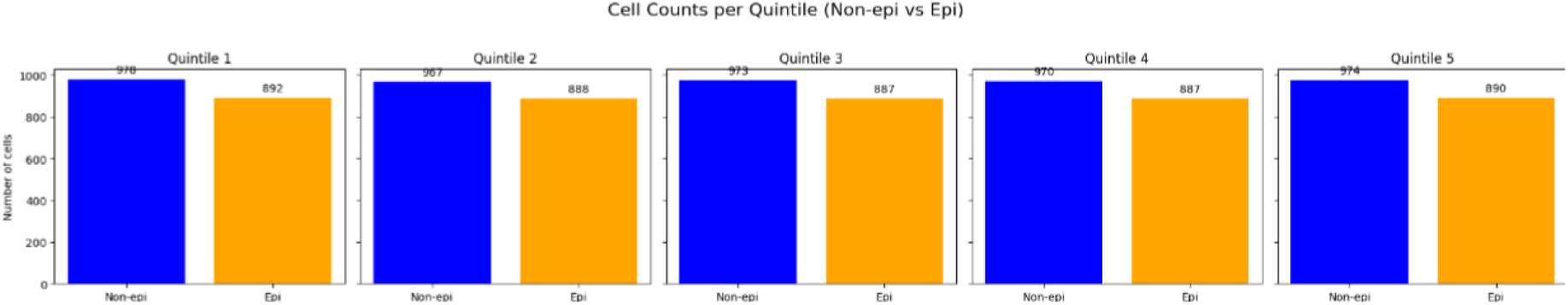
Cell counts across quintiles demonstrates no mismatch in the viable dataset to ensure model and metric ability to classify

Another concern was the potential for errors in the Danenberg study’s segmentation to be a confounding variable, preventing any biological interpretation. However, as demonstrated by the Random Forest Model and XGBoost models in addition to visual analyses, the vast majority of mis-segmented cells are above 150 pixels squared in terms of area, allowing them to be easily separated and have no impact on the rest of the dataset.

When performing feature importance in the XGBoost model (Figure 25 and 26), there was an interesting signal of a disproportionate reliance on minor axis length in 25, which resulted in an AUC of 0.7 and accuracy of 0.74. Exclusion of minor axis length from the model (26) had a negligible impact on overall accuracy (AUC: 0.68, Accuracy: 0.76), suggesting it may serve as a more easily accessible surrogate for broader morphological deviations, such as increased nuclear area, rather than an essential pillar for diagnosis.

**Figure 25:**
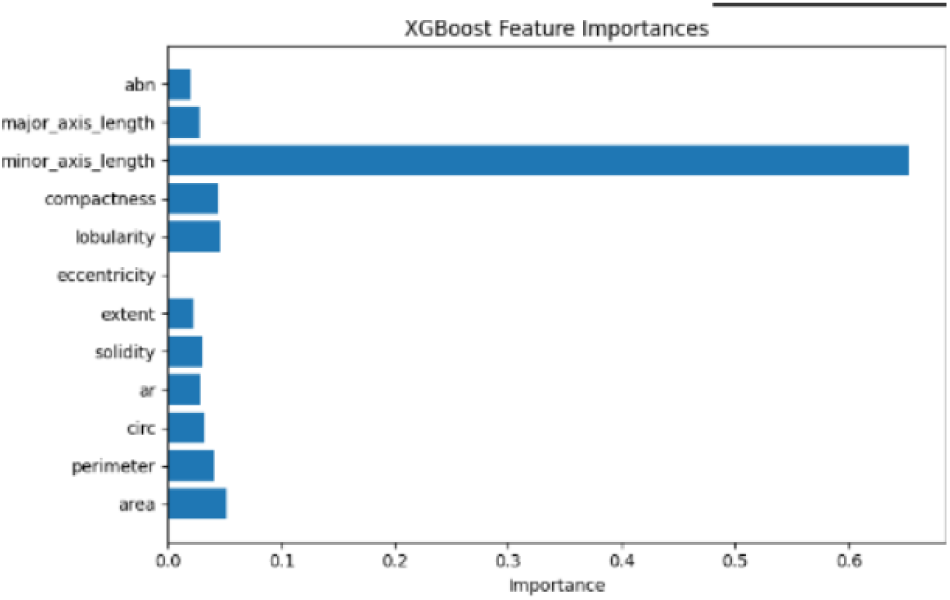
Importance of Minor Axis in XGBoost

**Figure 26:**
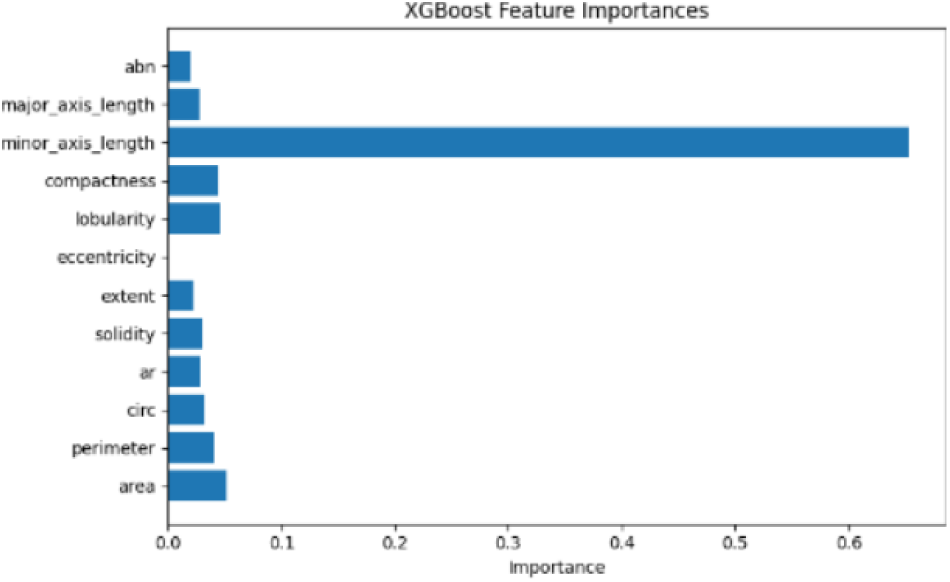
Feature importance without factoring in Minor Axis

To ensure that there was no relationship between cells of a lower area and the final abnormality, all metrics were plotted against area, and no statistically significant correlation was found as demonstrated in Figure 27. Cells with nuclear area below 20 pixels were visually inspected and excluded to minimize resolution artifacts. All small cells above the threshold of 20 pixels squared were inspected and found to be only affected by the biological signal rather than the potential limitations of IMC resolution.

**Figure 27:**
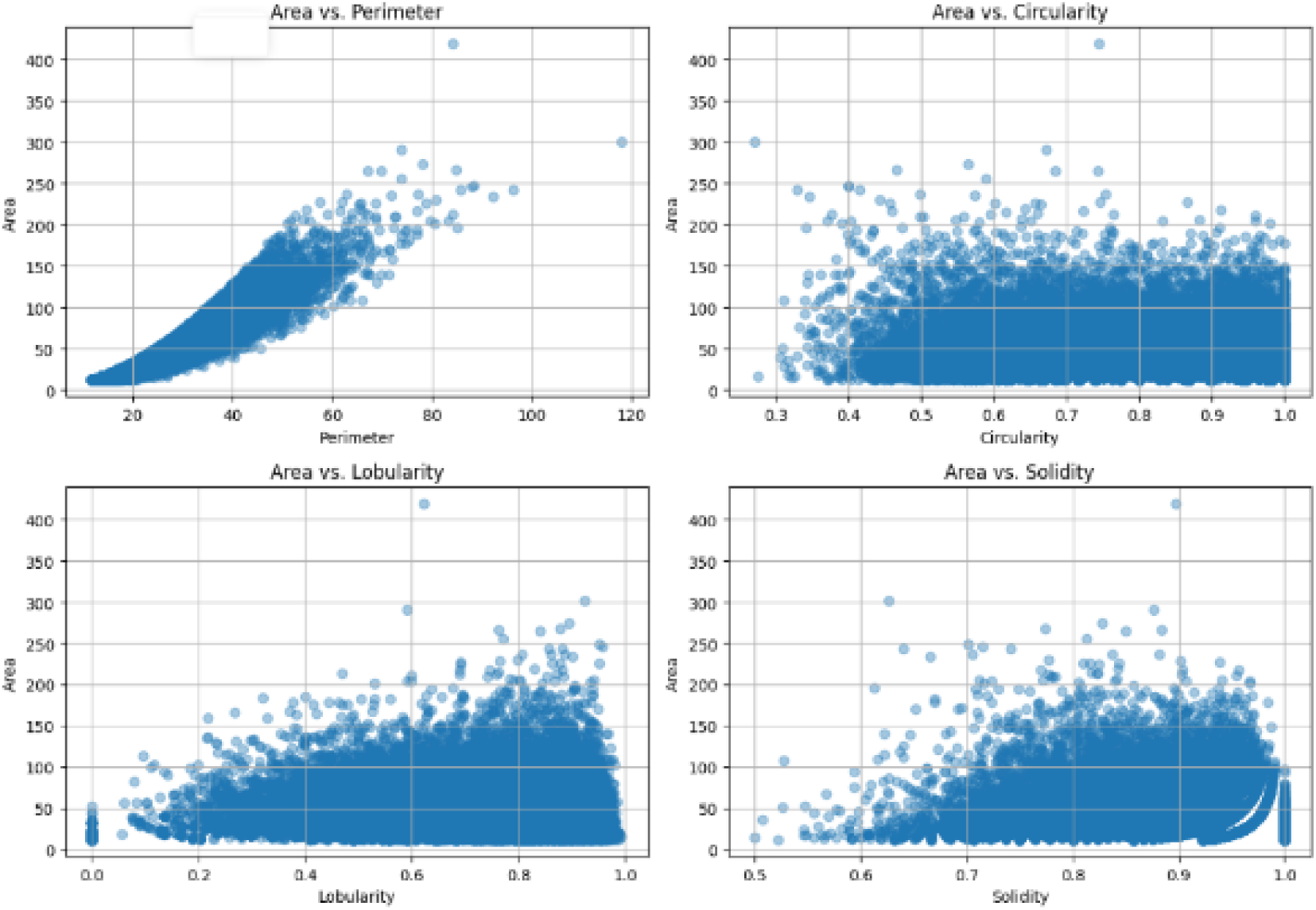
Scatter Plot of Area and Nuclear Metrics

In addition, a potential avenue for analysis is the inclusion of Zernike moments in 2D and 3D space, which are able to quantify irregularity irrespective of rotation or position potentially providing a solution to certain missegmentation errors. In testing, they provide a good baseline as a potential alternative to an abnormality score (AUC *<* 0.64); however, they were omitted due to the complexity of the calculations and the prioritization of efficiency.

**Figure 28:**
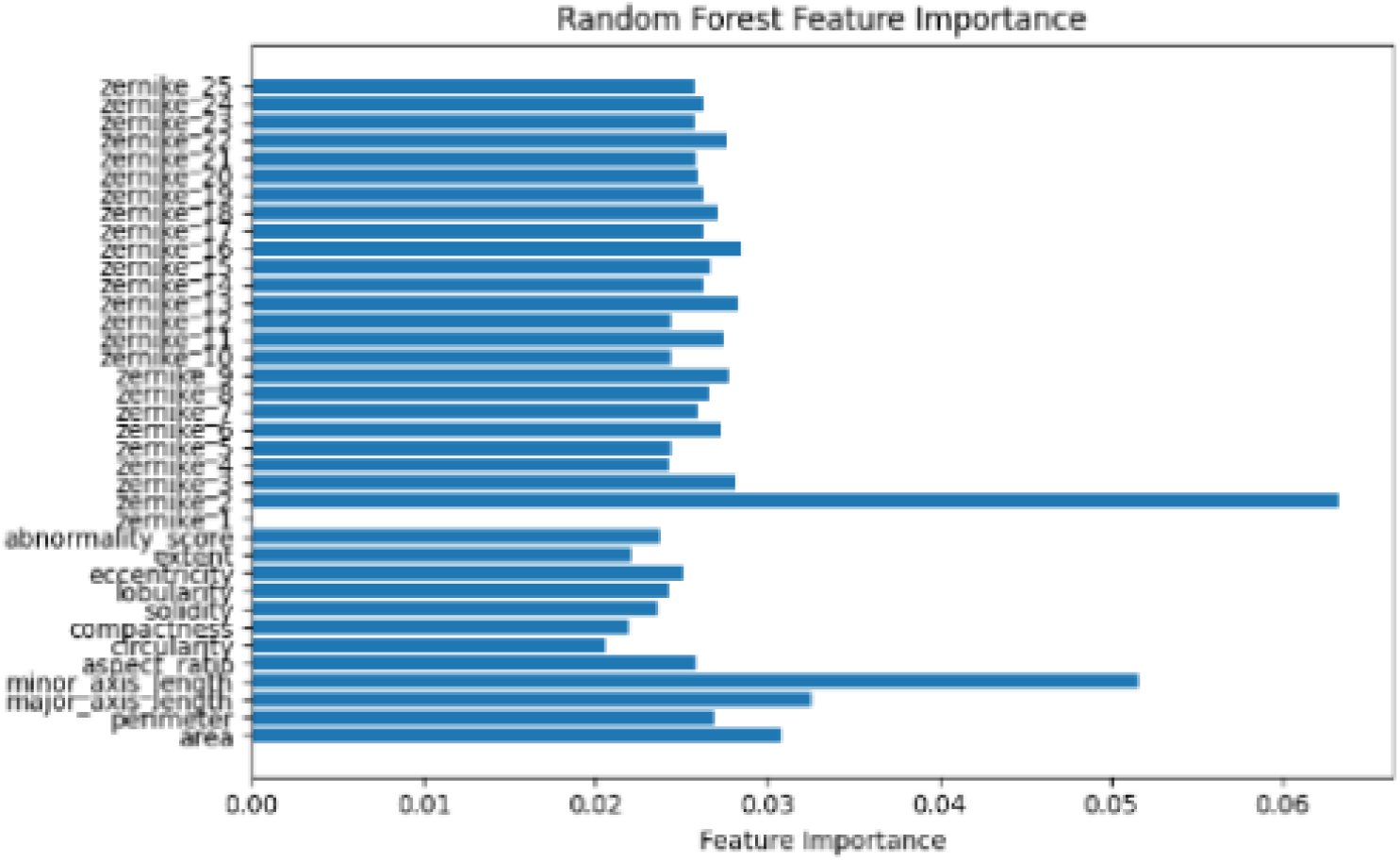
Random forest feature importance, including Zernike moments, demonstrates their ability to help classify in cancer above other established metrics

**Figure 29:**
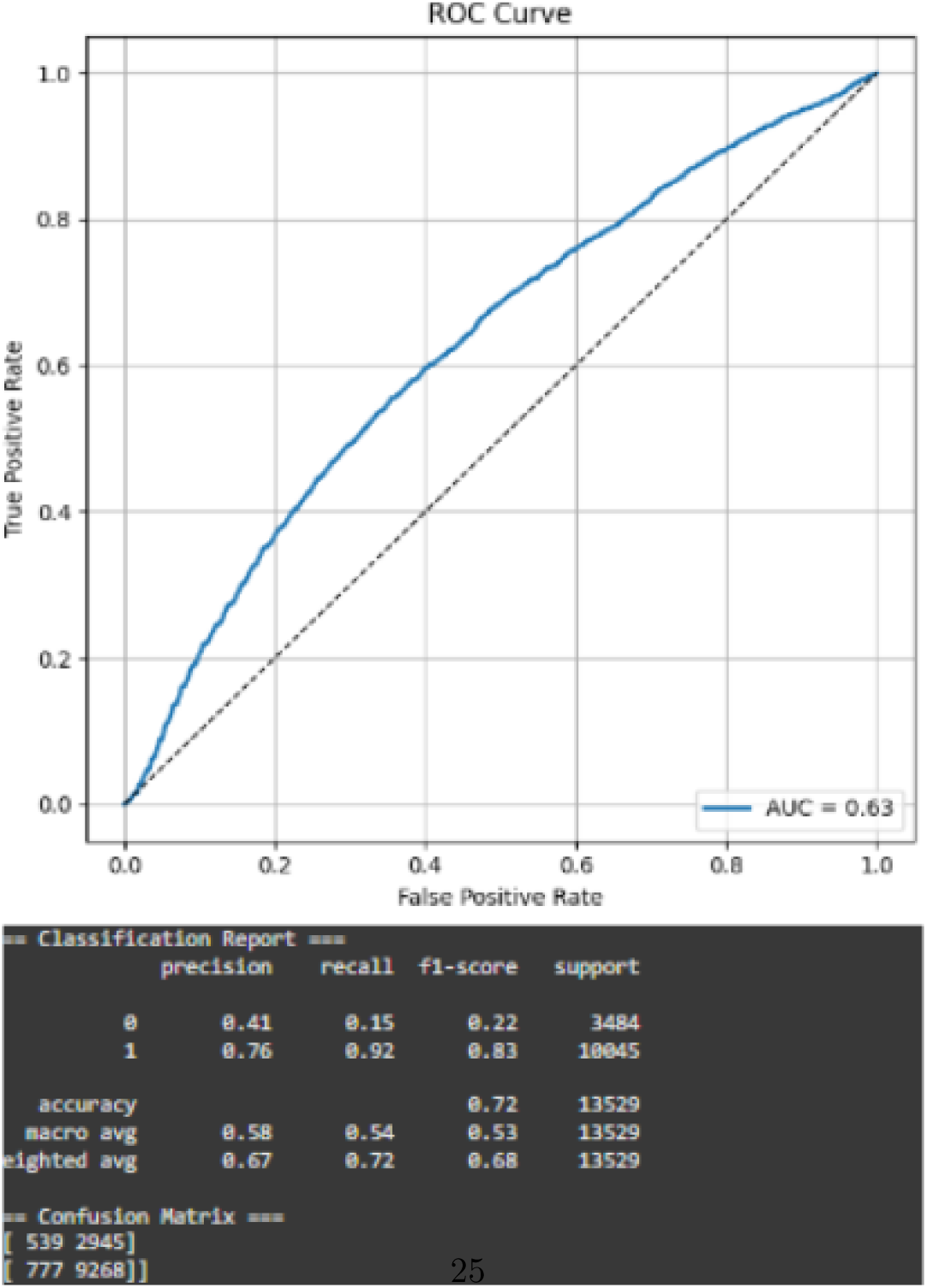
Zernike Moments Metrics achieve a high *<* 0.7 accuracy and *<* 0.64 AUC

While a preliminary analysis of markers compared to the abnormality score revealed important markers in IMC-based analysis, future studies can dive deeper into how marker analysis can be used in combination with nuclear metrics to better identify cancerous cells beyond simple regular and irregular scores. As seen below (Figure 30, and 31) preliminary analyses identify distinct populations of marker-expressing cells across both non-cancer and cancer markers, and a future step would be to find corresponding themes in nuclear morphological metric expression per cell type.

**Figure 30:**
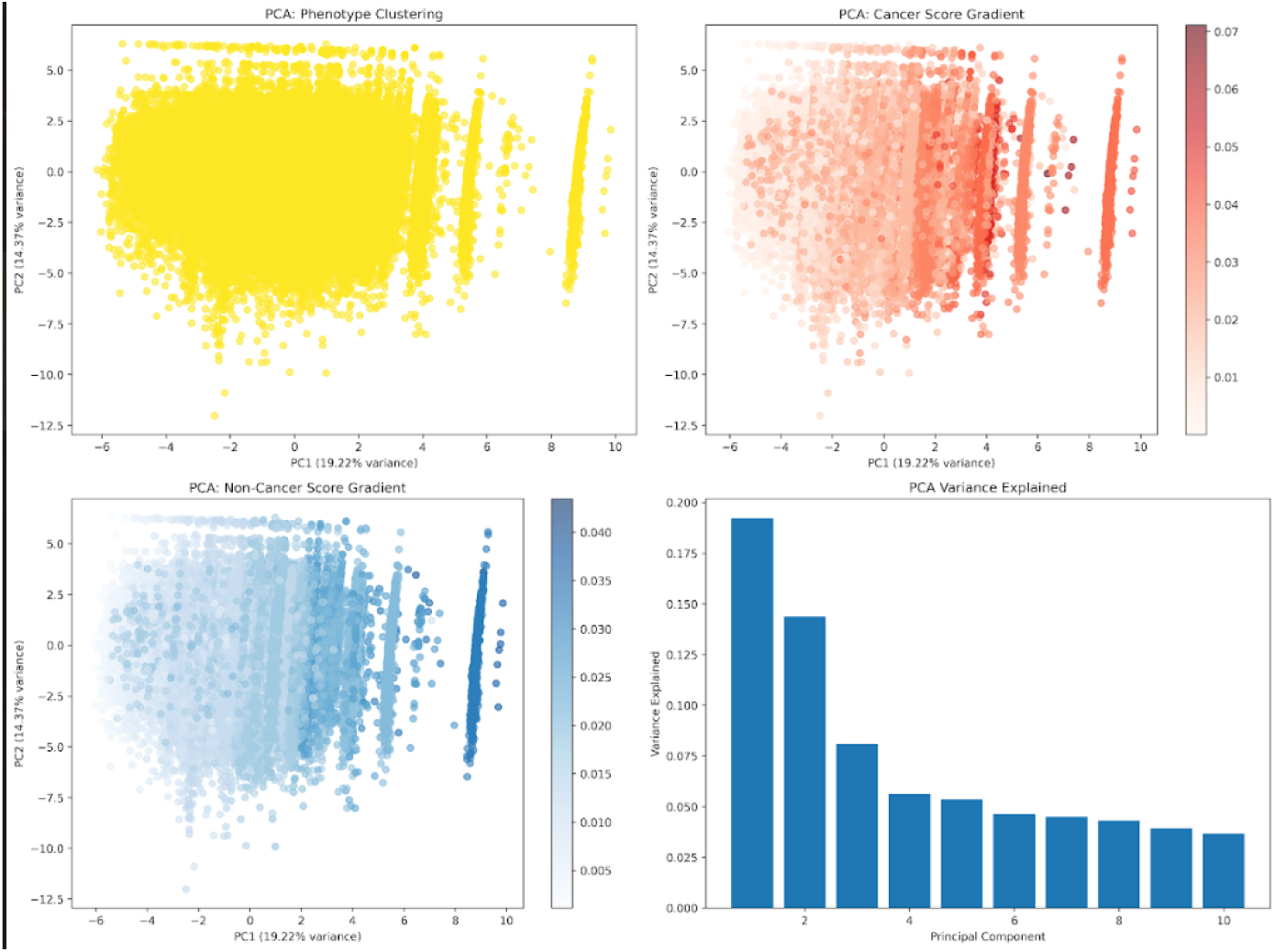
PCA marker importance displaying distinct clusters. Ranking of markers by their contribution to principal component variance reveals distinct populations.

**Figure 31:**
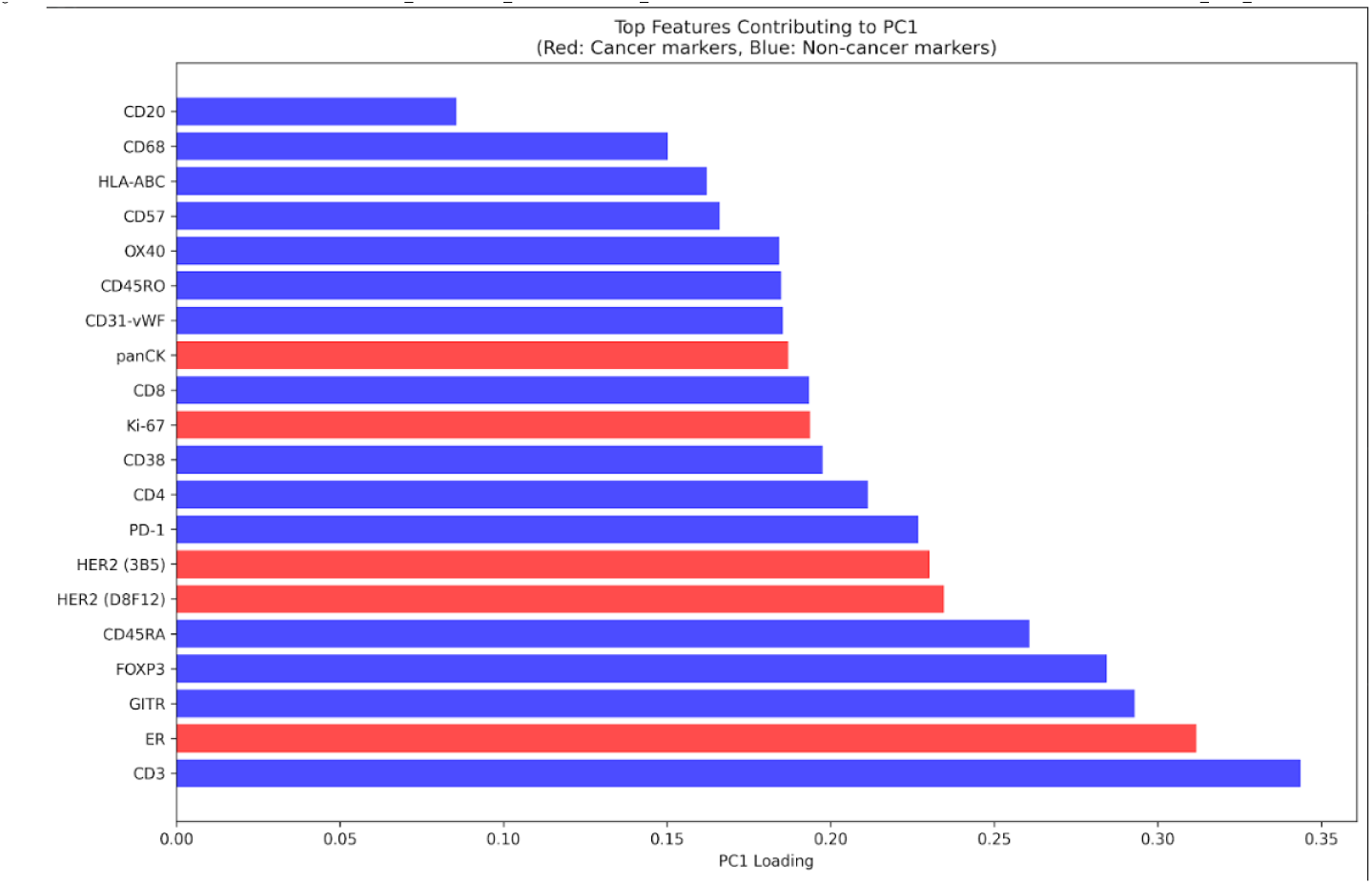
Marker signal distribution. Histogram or intensity spread of marker signals across cancer and non-cancer cells (as defined by the marker masks).

### 4.2 Conclusion

This study created new metrics to quantify nuclear morphology and was successfully able to separate epithelial cells by grouping cells by abnormality. This tool advances nuclear morphology metric analysis as a high-resolution, clinically translatable tool that sees nuclear deformation as a primary signal and insight into cancer state. The work displayed here lays the groundwork for a rapid, high-resolution classification of nuclear morphology across solid tumors using AI-powered segmentation. Future extensions of this workflow may utilize fully image-based virtual biopsies, eliminating the need for physical tissue extraction in diagnosis. In addition, by integrating nuclear shape metrics with genotypic data, this tool can be extended to infer underlying mutation status directly from morphology. This could streamline cancer subtyping and diagnosis in low-resource/income settings, allowing for longitudinal tracking of morphological drift within individual at-risk patients. It may also allow detection of early malignant transformation both in a laboratory setting and live in the operating room, detecting cells before clinical relapse or marker expression.

In summary, we developed a highly efficient, generalizable pipeline for analyzing nuclear morphology in breast cancer using imaging mass cytometry. Morphological deviations, across all quintiles, were strongly associated with malignancy, and this signal was confirmed across ML models and statistical tests. Our approach provides a clinical, translatable, and scalable process for cancer detection, with future work aimed at integrating spatial and omics data for even greater diagnostic potential across additional cancers. In addition, we discovered a minor axis and a cohesive cancer analysis metric that allows machine learning models to analyze nuclear masks and diagnose cancer in images.

## 5 Key Takeaways

1. Nuclear Morphology is a standalone tool for abnormal and cancerous cell identification
2. Abnormality quintiles reveal a cellular abnormality spectrum with the 5th quintile representing the most aggressive cancer subtypes that form a phenotype “tail”
3. Pipeline is generalizable across imaging data and cancer types due to reliance on cancer nuclei shape rather than specific markers
4. The abnormality score provides a compact, transparent biomarker for gate-based and AI cross-cancer analysis
5. Highly abnormal nuclei cluster spatially into regions of aggressive and passive subclones
6. This tool could be used for *rapid biopsy analysis* in aggressive cancer types like melanoma and breast cancer, cutting down time for cancer diagnosis by as much as 30 times
7. A GitHub comprised of nuclear analysis based tools was compiled for clinicians and researchers at: https://github.com/hamcoderfran/MM-IMC-Tool

## 6 Acknowledgements

Thanks to Dr. Kane Foster, Dr. Irene Ghobrial, my tutor Wesley Leong, RSI staff at MIT, CEE, Jason Koh, Shiv M. Gaglani, Ricardo Razon, Regeneron Pharmaceuticals Inc., and the Michael and Lori Milken Family Foundation

